# OPUS-TOMO: Towards Resolving Dynamics and Compositional Heterogeneities of Biomolecules with Cryo-Electron Tomography

**DOI:** 10.1101/2024.06.30.601442

**Authors:** Zhenwei Luo, Xiangru Chen, Qinghua Wang, Jianpeng Ma

**Affiliations:** Multiscale Research Institute of Complex Systems, Fudan University, Shanghai, 200433, China; Zhangjiang Fudan International Innovation Center, Fudan University, Shanghai, 201210, China; Shanghai AI Laboratory, Shanghai, 200030, China; Center for Biomolecular Innovation, Harcam Biomedicines, Shanghai, 201203, China

**Keywords:** Multiscale structural heterogeneity, high-resolution cryo-electron tomography, multi-body dynamics, ATP synthase dynamics, eukaryotic/prokaryotic ribosome translocation

## Abstract

Structural heterogeneity in biomolecules, arising from compositional and conformational variability, poses a major challenge for structure determination by cryo-electron tomography (cryo-ET). Here, we introduce OPUS-TOMO, a deep learning framework for resolving multiscale structural heterogeneity across the cryo-ET workflow. Our approach employs a convolutional encoder-decoder architecture and a rigid-body dynamics model to encode subtomograms into two separate low-dimensional latent spaces, realizing hierarchical modelling of compositional diversity and conformational dynamics. OPUS-TOMO adeptly resolved specie-level heterogeneity in template matching results and enabled sub-nanometer reconstructions for both *Chlamydomonas reinhardtii* ATP synthase dimer and *Schizosaccharomyces pombe* 80S ribosome, with an improvement of resolution up to 3 Å compared to expert-annotated datasets. OPUS-TOMO also resolved *in situ* dynamics for these biomolecules by reconstructing functionally relevant structural continua. The software is available at https://github.com/alncat/opusTOMO.

## Introduction

Recent advances in single-tilt axis holder^1^, cryo-focused ion beam (cryo-FIB) instruments^2,3^, and image processing algorithms^4–8^ have significantly propelled cryo-electron tomography (cryo-ET) towards sub-nanometer resolution^4^. Unlike cryo-electron microscopy (cryo-EM), cryo-ET preserves macromolecules in their native cellular environment^9,10^, enabling three-dimensional (3D) snapshots of “molecular sociology”^11^, and functional insights into *in situ* structures^12^. More importantly, cryo-ET could resolve *in situ* intermediate states for macromolecules that are impossible to obtain by conventional experiments. However, cryo-ET structural determination is bottlenecked by the structural heterogeneities.

Structural heterogeneities for biomolecules in cryo-ET arises at multiple scales including specie-level and molecule-level^13,14^. Initial particle picks from cryo-ET tomogram contain mixed biomolecular species due to multiple sources of high-level noises. Cryo-ET data have very low signal-to-noise ratios (SNRs) as a result of minimal-dose imaging^9,10,15^, missing-wedge artifacts from limited tilt angles of specimen holders^10,15^, defocus variations across the tilted images^9,15^ and crowded cellular environment^11^. Particle picking in cryo-ET cellular sample by template-based^16–18^ or template-free algorithms^19^ inevitably introduces large number of false positives, greatly limiting the resolution of reconstruction. Later particle sets purified by classification algorithms still exhibit significant dynamics and compositional heterogeneities on varying scales (inter-subunit or intra-subunit) due to the inherent functional flexibilities of biomolecules^20^. These heterogeneities, though biologically informative, limit the resolution of reconstruction and complicate biological discovery^10,11,21^. Therefore, a structural heterogeneity analysis framework that works across the whole cryo-ET pipeline is required to resolve multi-scale heterogeneities, either specie-level or molecule-level. It should be able to improve the resolutions of cryo-ET reconstructions^13^, and provide much-needed insights into the functionally relevant compositional changes and conformational dynamics.

Traditional approaches, which were implemented in pyTOM^7^, RELION^9^ and nextPYP^5^, rely on unsupervised machine learning algorithms and expert-designed strategies, such as hierarchical 3D classification^22^ to identify discrete states. Though it succeeds in resolving specie-level heterogeneity and revealing a number of important intermediate states via expert-designed strategies^23^, it falls short to reconstruct the structural continua which provide a panorama about the dynamical processes of biomolecules. Another line of research explicitly models the dominating low-frequency normal modes of the macromolecular dynamics (*e.g*., rigid-body subunit movements)^20^ while neglecting compositional variations. The multi-body refinement of RELION^24^ offers a low-dimensional, physically interpretable solution by parameterizing dynamics via rigid-body transformations for subunit. Similarly, 3DFlex in cryoSAPRC^25^, and Zernike/spherical harmonic-based approaches^26,27^ demonstrate the potential of neural networks and mathematical basis functions for modelling continuous deformation of biomolecules with cryo-EM data. Recent deep learning based methods in cryo-EM, such as cryoDRGN^28^ and OPUS-DSD^29^, leverage neural network based representation for 3D volume to model structural heterogeneities, inspiring cyro-ET adaptations like tomoDRGN^30^, and cryoDRGN-ET^31^. However, 3D volume representation models large-scale dynamics by a large number of voxel changes, thus being parameter-inefficient compared to low-dimensional physical model. In addition, these deep learning based frameworks lack demonstrations on resolving specie-level heterogeneities from template matching result, limiting their applications in the initial stages of cryo-ET pipeline.

In this paper, we report OPUS-TOMO, a deep learning framework that effectively resolves dynamics and compositional heterogeneities across the whole cryo-ET pipeline. OPUS-TOMO employs physical model to capture the inter-subunit dynamics, enabling efficient representation of large-scale movement. In OPUS-TOMO, two distinct latent spaces coexist: a dynamics latent space, paired with an MLP-based dynamics decoder to model inter-subunit movements, and a composition latent space, linked to a 3D convolutional neural network (3DCNN)-based composition decoder to capture compositional changes at all scales. This separation effectively decomposes the structural variations in different scales to different latent spaces. OPUS-TOMO revealed sub-nanometer structure for *C. reinhardtii* ATP synthase dimer, improving its resolution by 3 Å compared to expert-curated subtomogram set. On diverse real-world datasets, it achieved resolution gains of up to 4.5 Å, often reaching the Nyquist limit, while resolving dynamic molecular transitions *in situ*. Specifically, OPUS-TOMO characterizes the dynamics of ATP synthase, i.e., the F_1_ Head accompanies the 120° rotation of central stalk through a rotation of ∼30° during the transition between rotatory states^32^. OPUS-TOMO reveals the function of eEF2 as an anchoring point during tRNA translocation by reconstructing an important translocation-intermediate state with two tRNAs trapped in hybrid the pe/E and ap/P positions^33^, illustrating the interplay between function and dynamics in biomolecules. By automating and scaling heterogeneity analysis, OPUS-TOMO can facilitate cryo-ET structure determination and yield essential biological insights at molecular level for biomolecules by reconstructing continuous intermediate states *in situ*.

## Results

### Neural Architecture of OPUS-TOMO

Template matching and subtomogram averaging (STA) represent two essential steps in cryo-ET structure determination. Specifically, template matching finds the location and orientation of a target macromolecule inside a tomogram, whereas subtomogram averaging yields a reconstruction by determining translations and orientations of subtomograms with respect to (w.r.t) the subtomogram average. The architecture of OPUS-TOMO is outline schematically in **Fig. 1a**. The input of OPUS-TOMO is from either the template matching or subtomogram averaging results. The 3D subtomogram is first aligned to the canonical orientation of a template or a subtomogram average by inversely rotating according to its estimated pose. The encoder takes the aligned subtomogram, and estimates the distributions of its composition latent code *z* and dynamics latent code *z*_*d*_. Specifically, it estimates the mean *z* ∈ ℝ^*n*^ and standard variation 𝜎 ∈ ℝ^*n*^ for each latent code^34^ (**Fig. 1a**). Next, the latent codes are sampled according to the estimated distributions and supplied to the corresponding decoders. The sampled composition latent code is supplied to the composition decoder to reconstruct a 3D volume *V*(***x***) (**Fig. 1a**), which is represented by a discrete grid of voxels. Meanwhile, the sampled dynamics latent code is supplied to the dynamics decoder to estimate a deformation field (**Fig. 1a**), which mainly serves to correct deformation of complex originating from the relative movements between subunits. Finally, the spatial transformer^35^ is leveraged to produce a 3D reconstruction by applying the deformation field with the supplied pose onto the 3D volume *V*(***x***) (**Fig. 1a**). The 3D reconstruction is transformed into a subtomogram with its corresponding 3D-contrast transfer function (3DCTF) following the differentiable model defined in the subsection of **Subtomogram Formation Model** in **Methods**.

**Figure 1.**
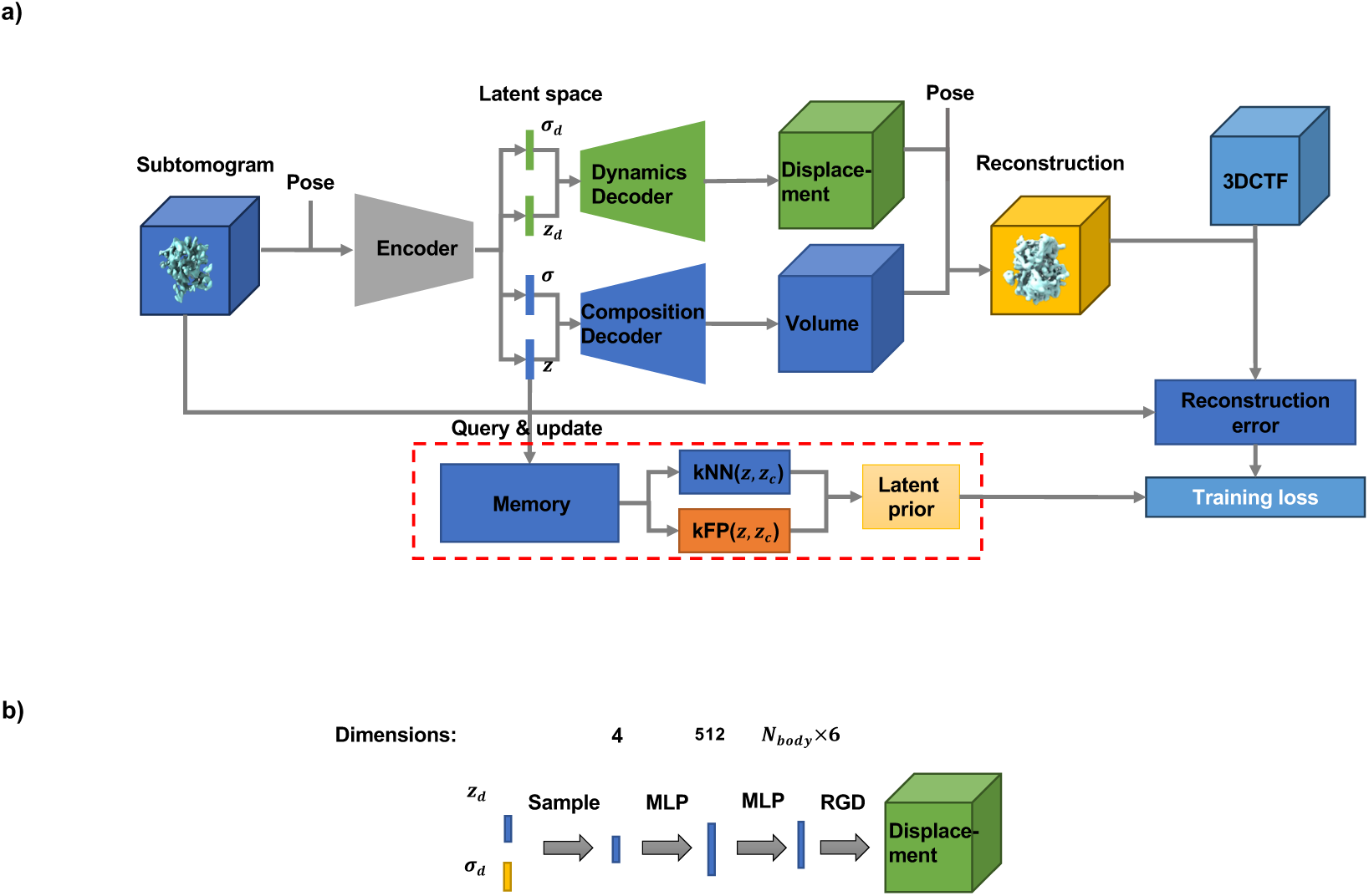
OPUS-TOMO architecture. **a)**. **Overall architecture.** Pose refers to the projection direction of the subtomogram with respect to the consensus subtomogram average. 3DCTF refers to its 3D contrast transfer function. *z* is the mean vector of composition latent code. 𝜎 is the standard deviation vector of composition latent code. *z*_*d*_ is the mean vector of dynamics latent code. 𝜎_*d*_ is the standard deviation vector of dynamics latent code. kNN(*z*, *z*_*d*_) refers to the K nearest neighbor of the concatenation of *z* and *z*_*d*_. kFP(*z*, *z*_*d*_) refers to the K farthest points of the concatenation of *z* and *z*_*d*_. For better visualization, the link between the dynamics latent code and the memory is omitted. The standard Gaussian prior for latent distribution as well as the smoothness and sparseness priors for the 3D volume are also omitted. **b)**. **Architecture of the dynamics decoder of OPUS-TOMO.** This diagram shows the dynamics decoder that translates the latent encoding *z*_*d*_ into a deformation field. MLP denotes the multi-layer perceptron. RGD denotes the rigid-body dynamics model that transforms the predicted rigid-body movement parameters into the 3D deformation field. All MLPs except the last one are with LeakyReLU nonlinearities having a negative slope of 0.2.

The neural architecture of OPUS-TOMO can be trained end-to-end by reconstructing each input subtomogram and minimizing the squared reconstruction errors between the input and the output (**Fig. 1a**). During training, to encourage the smoothness of latent spaces, the composition and dynamics latent codes are regularized by the structural disentanglement prior^29^ from OPUS-DSD and Kullback-Leibler (KL) divergence w.r.t standard Gaussian^34,36^ (**Fig. 1a**). The encoder and composition decoder utilizes the 3D convolutional networks from OPUS-DSD^29^. The dynamics decoder consists of a series of MLPs and nonlinearities (**Fig. 1b**). The dynamics latent code is transformed into a 512-dimensional representation by an MLP followed by a LeakyReLU nonlinearity. The 512-dimensional representation is projected to a *N*_body_ × 6-dimensional vector by another MLP, where *N*_body_denotes the number of rigid bodies defined for the macromolecule. This *N*_body_ × 6-dimensional vector is then reshaped into *N*_body_ vectors, each of 6 dimensions, which define the rotation and translation parameters for each subunit’s rigid-body dynamics (**Fig. 1b**). The first three dimensions of the 6-dimensional vector define the rotation using a quaternion representation, while the last three dimensions defines the translation of the subunit. The construction of the deformation field using the rotation and translation parameters is detailed in the **Deformation field** subsection in **Methods**.

### High-resolution Reconstruction of Mitochondrial ATP synthase Dimer from Tomograms of *Chlamydomonas* (*C*.) *reinhardtii*

ATP synthase is a protein complex embedded in the crista membrane, and serves to synthesize the ATP using the proton gradient across the membrane. Recently, the structure of ATP synthase dimer in *Polytomella* cells was determined to a sub-nanomter resolution (8.6 Å) using *in situ* cryo-ET^23^. Meanwhile, the structure of ATP synthase in *C. reinhardtii* cells was also revealed by cryo-ET^37^. Though the structures for peripheral stalk and F_1_ head domain were determined to sub-nanometer resolutions by focused refinements, the structure for *C. reinhardtii*ATP synthase dimer in sub-nanometer resolution is not reported yet. Moreover, both teams resolve the ATP synthase by iterative rounds of 3D classifications on template matching results, which is extremely labor-intensive and time-consuming. Here, we demonstrated the capability of OPUS-TOMO for enabling sub-nanometer cryo-ET reconstruction from template matching results using only two rounds of filtering, greatly accelerating the tomography workflow.

261 tomograms for *C. reinhardtii* mitochondria were reconstructed by WARP^6^ using the tilt series and their corresponding parameters of tilt series alignment deposited in EMPIAR-11830^37^. Using template matching in pyTOM^16^ with a published *Polytomella* ATP synthase map^23^ (EMDB accession code: EMD-50000) low-pass filtered to 20Å as the template, we picked a total of 783,000 subtomograms (3000 per tomogram) and exported them in 3.92 Å by WARP. The first round of OPUS-TOMO training served to filter out the large number of membrane false positives in the template matching set. During training, the dynamics decoder treats the whole structure as a single rigid-body to correct the pose from template matching. Firstly, we used subtomograms from 62 randomly chosen tomograms to train an initial model for filtering membranes. The UMAP visualization of the 12-dimensional composition latent space learned by OPUS-TOMO shows distinct clusters, which we clustered into 20 classes by KMeans algorithm, and reconstructed 3D structure for each class by supplying cluster centers into the trained composition decoder of OPUS-TOMO (**Extended Data Fig. 1a**). Class 13 (10,709 subtomograms) shows complete densities for ATP synthase dimer, and Class 15 (43,146 subtomograms) shows weak densities resemble ATP synthase dimer, while remaining classes are membranes and unknown particles (**Extended Data Fig. 1a**). Subtomograms from Classes 13 and 15 (29% of subtomograms from the 62 tomograms) are kept for further processing. The subtomograms from remaining tomograms are filtered by training OPUS-TOMO with the pretrained initial model and KMeans clustering in latent space following the same criteria. The first round of filtering resulted in 220,291 subtomgrams (28% of subtomograms from all tomograms).

The second round of filtering was performed by training OPUS-TOMO on the 220,291 subtomograms with orientations from template matching. The UMAP visualization of the 12-dimensional latent space learned by OPUS-TOMO shows subtomograms for the ATP synthase as separated clusters compared to non-ATP synthase species (**Extended Data Fig. 1b**). Impressively, Class 0 and Class 1 show ATP synthase tetramer, while Class 3 shows ATP synthase hexamer, indicating the formation of higher order ATP synthase oligomer *in situ* (**Fig. 2a**). Moreover, those oligomers are interlocked through their ATP synthase-associated protein (ASA) subunits (red label in **Fig. 2a**) as it revealed in *Polytomella* ATP synthase structure *in situ*^23^. All 33,338 subtomograms from Classes 0∼3 were imported into M^4^ for refinement by focusing on the central ATP synthase dimer and enforcing *C*_2_ symmetry. Classes 0∼3 yielded a subtomogram average with a resolution of 8.9 Å after multiple rounds of 2D image warp, CTF and particle pose refinement in M^4^ (**Fig. 2b**). To validate the existence of high-order oligomers of ATP synthase dimer, we added 14,352 subtomograms centering at the up ATP synthase dimers in Class 0 and Class 3 to the previous set. The expanded set contained 47,690 subtomograms, which yielded a subtomgram average with the same resolution 8.9 Å but showing improved local resolutions under the same refinement settings, corroborating the existence of high-order oligomers of ATP synthase dimer identified by OPUS-TOMO (**Fig. 2b**). Compared to the 6.8 Å *Polytomella* ATP synthase structure (EMD accession code: EMD-50001)^23^, the subtomogram averages obtained by OPUS-TOMO show helical structures with similar resolutions but lacks “bridge” dimer linkage between the first subunit of ATP synthase-associated protein (ASA1) subunits in the upper part of the peripheral stalk^23^, which is a significant difference between the *C. reinhardtii* and the *Polytomella* ATP synthase dimer *in situ*^37^ (**Fig. 2c**).

**Figure 2.**
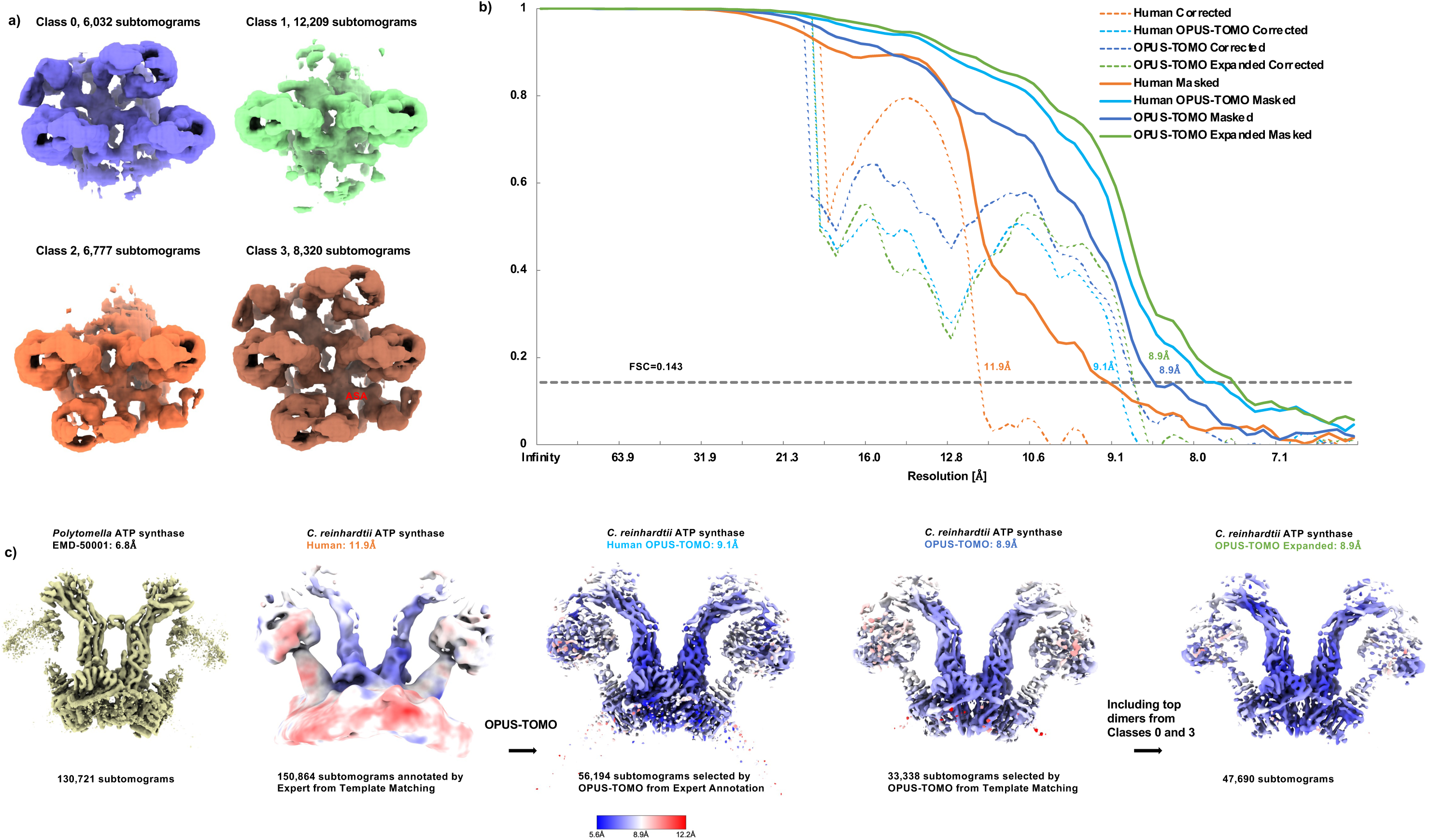
Oligomers and reconstructions of the *C. reinhardtii* ATP synthase dimer *in situ* (data from EMPIAR-11830). **a).** Different oligomers of ATP synthase dimer revealed by OPUS-TOMO (**Extended Data Fig. 1b**). Dimers are interlocked through their ASA subunits (red label). **b)**. Comparison between the gold standard Fourier shell correlations (FSCs) of the subtomogram averages obtained by different datasets. The FSC curve is named using a combination of prefix and suffix. The suffix represents the type FSC, where “Corrected” and “Masked”, represent corrected FSC and masked FSC, respectively. The prefix specifies the dataset for reconstructing the corresponding subtomogram average. “Human” represents the set of subtomograms from deposited expert annotations^37^. “Human OPUS-TOMO” represents the set of subtomograms from Classes 3∼13 classified by OPUS-TOMO on “Human” set (**Extended Data Fig. 1c**). “OPUS-TOMO” represents the set of subtomograms from Classes 0∼3 classified by OPUS-TOMO in the second round of filtering on template matching results (**Extended Data Fig. 1b**). “OPUS-TOMO Expanded” represents the set of subtomograms composed by combining the center dimers in Classes 0∼3 with top dimers from Class 0 and Class 3 (**Fig. 1a)**. The FSC curves and the resolutions at FSC=0.143 are colored by origins of datasets. The color scheme is consistent throughout this figure. **c)**. Subtomogram averages of ATP synthase dimer obtained by different datasets. The number of subtomograms in each dataset is annotated below. Density maps except “Human” are contoured at 5.5𝜎 level, and colored by local resolutions except the *Polytomella* ATP synthase. The density map for “Human” is contoured at 4𝜎 level to show similar structure.

The ATP synthase set for *C. reinhardtii* annotated by Ron et. al.^37^ served as a baseline to compare the performance of OPUS-TOMO to tomography workflow based on human expert. The expert-annotated ATP synthase set for *C. reinhardtii* which contains 150,864 subtomograms yielded an average with a resolution of 11.9 Å after refinements in M using the same settings (**Fig. 2b**). The density map from the expert-annotated set failed to reveal high-resolution helical structures compare to the set produced by OPUS-TOMO (**Fig. 2c**). To further demonstrate the capability of OPUS-TOMO, we reconstructed the subtomograms from the expert-annotated set using WARP, and trained OPUS-TOMO on them. In the UMAP visualization of the 12-dimensional composition latent space learned by OPUS-TOMO, which we clustered into 30 classes by KMeans algorithm, Classes 3∼13 representing ATP synthase oligomers form a distinct cluster (**Extended Data Fig. 1c**). Further refinements on Classes 3∼13 (56,194 subtomograms) yielded an ATP synthase dimer of resolution 9.1 Å (**Fig. 2b**), and revealed the high-resolution helical structures which are similar to EMD-50001 (**Fig. 2c**).

In the reconstruction of ATP synthase dimer, we demonstrated that heterogeneity analysis by OPUS-TOMO was essential for obtaining high-resolution structures. OPUS-TOMO significantly outperformed human expert by producing purified subtomogram sets which achieved resolutions at 8.9 Å from template matching result, and improving the final resolution of ATP synthase dimer by 3 Å.

### Rotary States and Dynamics of Mitochondrial ATP synthase Dimer from Tomograms of *C. reinhardtii*

ATP synthase is a protein complex consisting of a proton-translocating membrane domain, F_o_, attached via central and peripheral stalks to the catalytic domain F_1_. The rotation of central stalk and the associated *c*-ring being powered by the transmembrane proton-motive force induces conformational changes in F_1_ head that result in ADP phosphorylation^32,38^. Those large-scale functional conformational changes, especially in F_1_ heads and central stalks, lead to broken densities in those regions in the previously determined structures of ATP synthase dimer. Therefore, we continue to reveal the dynamics and compositional heterogeneity which are crucial for understanding the ATP synthesis *in situ* by training OPUS-TOMO on the M-refined OPUS-TOMO expanded set. During training, the dynamics decoder leveraged a two-subunit model to capture the relative displacements between two monomers, in which each subunit is centered at the center of mass (COM) of the F_1_ Head of each monomer.

The 360° rotation of the central stalk with a elementary step of 120° rotation divides the ATP synthase into three primary rotary states^32^. OPUS-TOMO can easily reveal them by clustering the learned composition latent space and reconstructing the 3D structure at each cluster center using the trained composition decoder (**Extended Data Fig. 2a**). For illustration, we compared Class 7, Class 16 and Class 19, which were located in separated clusters in latent space, to the corresponding rotary states of *Polytomella* ATP synthase (**Fig. 3a** and **Extended Data Fig. 2a**). The three primary rotary states show different orientations of the central stalk with a 120° separation. The top of central stalk of Class 7 orients left, the top of central stalk of Class 16 points to middle, while the top of central stalk of Class 19 orients right, which are evident in separate and superposition view (**Fig. 3a∼b**). In the sectional view along the Y axis at the ADP binding sites, the top of central stalk extends to different 𝛽 subunits in this resolution, and evenly partitions the circle of F_1_ head in superposition view, thus confirming the 120° separation of central stalk in different states (**Fig. 3c∼d**). The identities of these states are supported by the overlays of the corresponding atomic structures of ATP synthase (**Fig. 3a**) and high-resolution density maps (**Extended Data Fig. 2b**). OPUS-TOMO can also reveal the rotary substates for ATP synthase. For example, superposition of the density maps for Class 18 and Class 19 reveals the Class 19 as a rotary substate for state 3 (**Fig. 3e**). Though their central stalks adopt the same orientation, the F_1_ Head of Class 18 rotates by a small degree and makes the 𝛽 subunit near peripheral stalk bind with ASA1 at this resolution (**Fig. 3e∼f**). The existence of this substate indicates the possible role of ASA1 in mediating the rotation of F_1_ Head.

**Figure 3.**
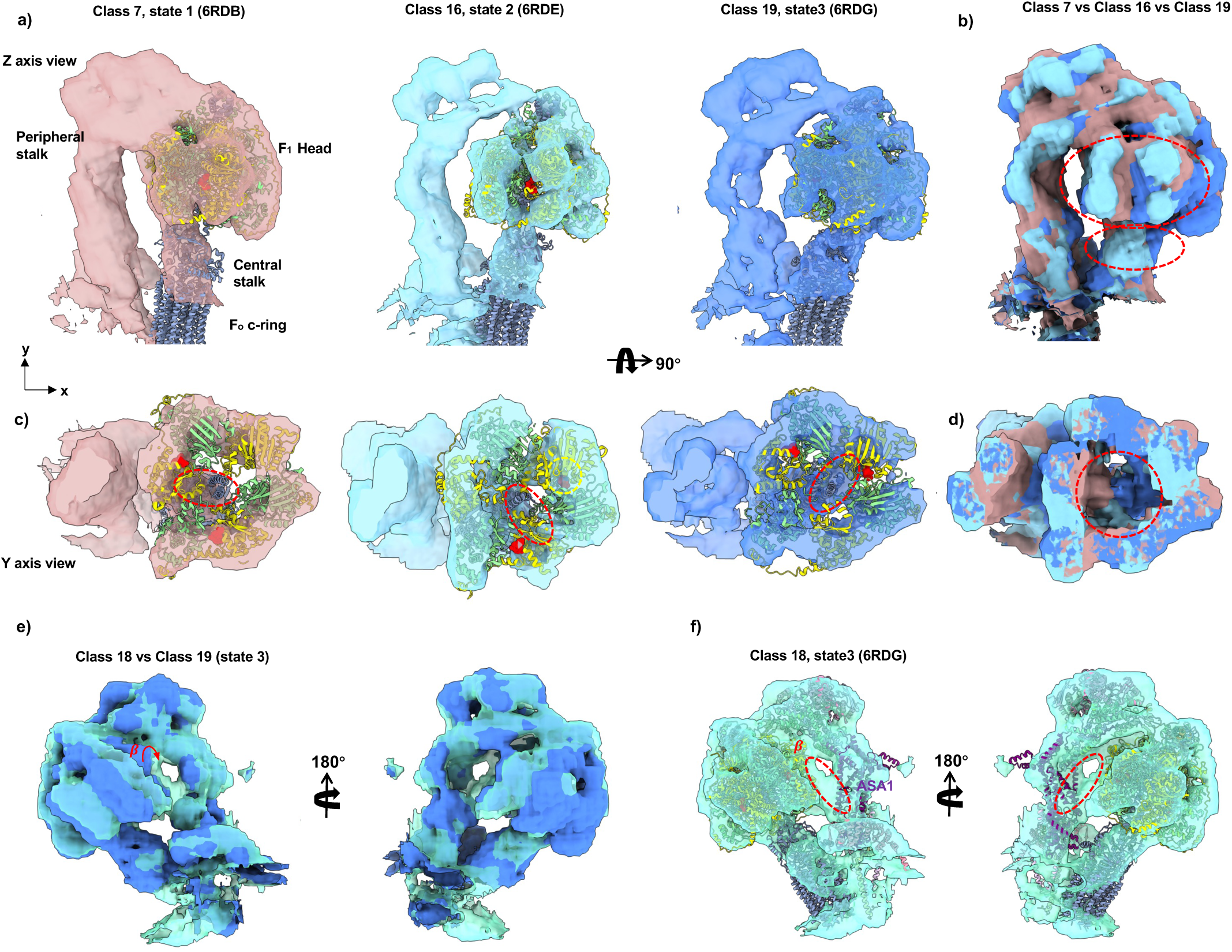
Rotary states of the *C. reinhardtii* ATP synthase *in situ*. **a)**. Three primary rotary states (1, 2, and 3) of ATP synthase reconstructed by OPUS-TOMO at different class centroids (**Extended Data Fig. 2a**). The correspondence between the class from OPUS-TOMO and the rotary state is established by fitting the structure of rotary state (PDB code in parentheses) into the density map for this class. The structure of central stalk (dark blue) overlays with the density of central stalk, verifying the correctness of the orientation of central stalk in density map for different rotary states. The density map of F_1_ Head encloses 𝛼 subunits (light green) and 𝛽 subunits (yellow). **b)**. Superposition of density maps exposes that the top of central stalk (small red dashed ellipse) points to left (Class 7, brown), middle (Class 16, light blue) and right (Class 19, blue) in different states. The positions of the 𝛼 and 𝛽 subunits also vary in different states (large red dashed ellipse). **c)**. Section through F1 at the level of bound ADP (red spheres). The top of central stalk extends to different 𝛽 subunit in this resolution (red dashed ellipses). **d)**. Superposition of density maps evenly partitions the F_1_ Head into three parts (red dashed circle), confirming the 120° degree rotation of the central stalk in these states. **e)**. Rotary substates of state 3. Superposition of Class 18 (transparent green) and Class 19 (solid blue) shows the relative displacement of 𝛽 subunit (red text and arrow) between different rotary substates of state 3. **f)**. 𝛽 subunit (red text) in Class 18 binding (red dashed ellipse) with the ASA1 subunit (purple text), indicating that ASA1 possibly mediates the rotation of F_1_ Head.

Except reconstructing different rotary states, a more intriguing problem is revealing the continuum for the elementary 120° rotation of the central stalk, which requires reconstructing multiple substates to link primary states. They were revealed by hierarchical focused classification in previous studies^23,32^. OPUS-TOMO can directly reconstruct such a continuum by traversing the principal component (PC) of the learned composition latent space. Specifically, the negative end of PC11 (−2.8PC11) is found to be the state 2, the middle point of PC11 (0.8PC11) correspond to state 2A (rotary substate of state 2), while the positive end of PC11 (3.6PC11) is found to be state 1A (rotary substate of state 1)^32^, thus reconstructing the transition from state 2 to state 1A via state 2A (**Fig. 4a)**. The identities of −2.8PC11 and 3.6PC11 were verified by the fitting of the corresponding atomic structures of ATP synthase (**Fig. 4a**) and high-resolution density maps (**Extended Data Fig. 3a**). The superposition of density maps from different states and the corresponding fitted atomic structures reveals a set of conformational changes in different subunits during transition (**Fig. 4b**). First, the OSCP domain and the nearby ASA4 subunit moves leftwards, charactering the movement of OSCP and peripheral stalk (**Fig. 4c**). Second, the 𝛾 subunit rotates by 120° through a 60° rotation at state 2A, characterizing the movement of central stalk (**Fig. 4d**). Third, the 𝛽 subunits rotate by ∼30° through a ∼15° rotation at state 2A (**Fig. 4e∼g**). The rotation of 𝛽 subunit stops until the nearby 𝛼 subunit contacts with ASA4 (**Fig. 4a, 4e**), which indicates the possible role of ASA4 in mediating the rotation of F_1_ Head. In summary, this path reconstructed by OPUS-TOMO characterizes the 30° rotation of F_1_ Head accompanying the 120° rotation of central stalk and is consistent with the previous cryo-EM studies^32^ (**Supplementary Video 1**).

**Figure 4.**
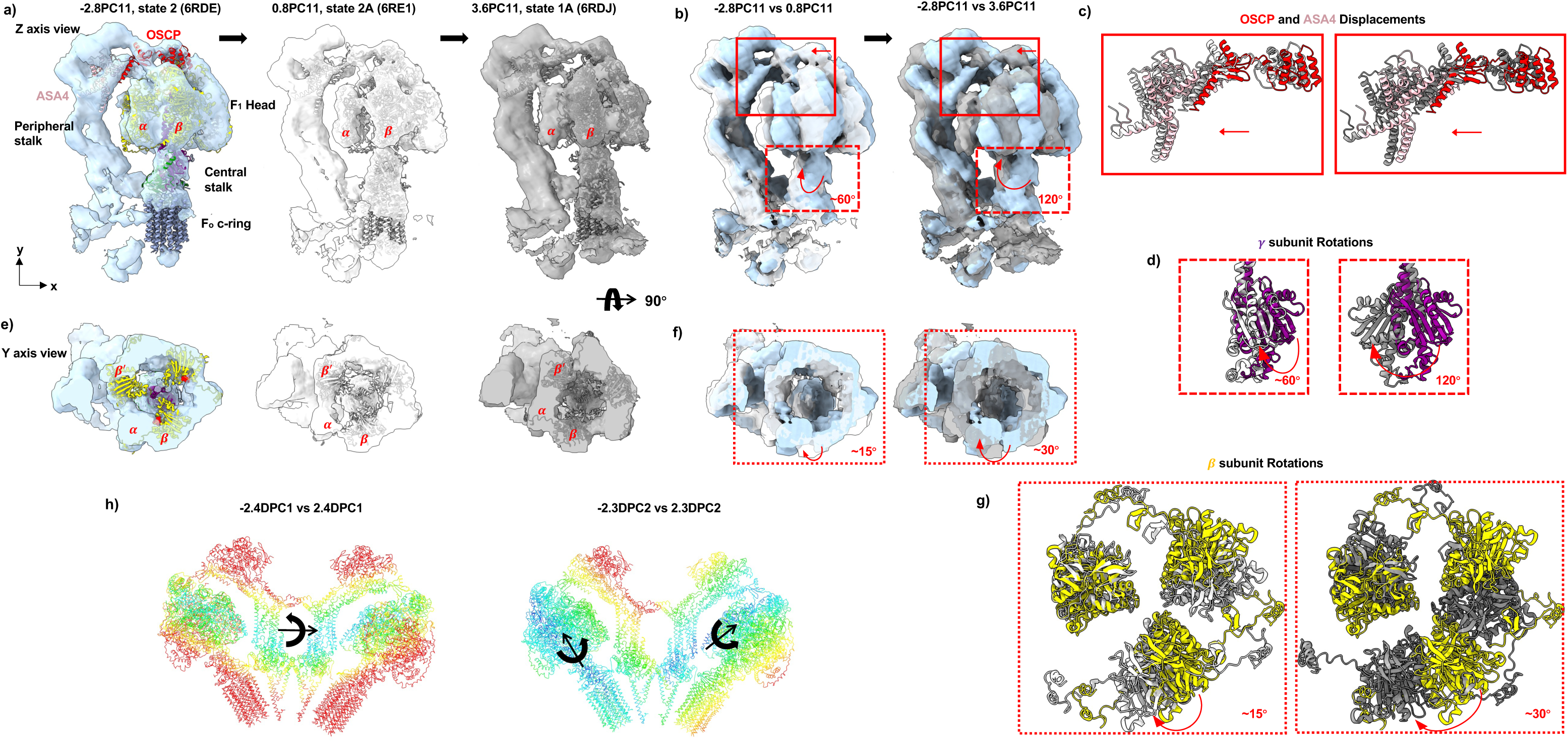
Dynamics of the *C. reinhardtii* ATP synthase *in situ*. **a)**. Transition between rotary substates revealed by progressing in the direction of PC11 in OPUS-TOMO’s composition latent space. The location of latent code where the density map is reconstructed by the composition decoder of OPUS-TOMO is specified as xPC11. The correspondence between xPC11 and rotary substate is established by fitting the atomic structure of the rotary substate (PDB code in parentheses) into the map at xPC11. The 𝛼 and 𝛽 subunits (red text) rotate towards peripheral stalk during the transition. **b)**. Superpositions of density maps show increasingly large concerted displacement of the OSCP and ASA4 subunits (red boxes) from −2.8PC11 (blue) to 3.6PC11 (grey) through 0.8PC11 (light grey) in the direction given by red arrows. Meanwhile, the central stalk (red dashed boxes) rotates by ∼60° at 0.8PC11, and 120° at 3.6PC11 in the direction given by red arrows. **c)**. Closeup view of the displacement between the OSCP and ASA4 subunits fitted to densities of different rotary states. The OSCP and ASA4 fitted to the starting state (−2.8PC11, state 2) are shown in red and pink, respectively. All subunits fitted to other states are shown in the colors of the corresponding density maps. **d)**. Closeup view of the rotation of 𝛾 subunit fitted to densities of different rotary states. The 𝛾 subunit fitted to the starting state is shown in purple. **e)**. Section through F_1_ at the level of bound ADP (red spheres). 𝛽′ subunit rotates away from the peripheral stalk, confirming the global rotation of F_1_ Head. **f)**. Superpositions of density maps show the rotation of F_1_ Head (red dotted boxes) from −2.8PC11 (blue) to 3.6PC11 (grey) through 0.8PC11 (light grey) in the direction given by red arrows. **g)**. Closeup view of the rotation between the three 𝛽 subunits fitted to densities of different rotary states. From −2.8PC11 (state2, yellow) to 0.8PC11 (state 2A, light grey), the 𝛽 subunits rotate by ∼15°. From −2.8PC11 (state2, yellow) to 3.6PC11 (state 1A, grey), the 𝛽 subunits rotate by ∼30°. **h)**. Color-coded displacement maps reveal two orthogonal inter-monomer movements along different DPCs of OPUS-TOMO’s dynamics latent space. Residues in the structures are colored by their displacement distances along the corresponding DPCs (**Extended Data Fig. 3b∼c**), using rainbow palette in ChimeraX^57^ with a scale from 0 to 15 Å, where red indicates larger displacements and blue suggests smaller displacements. Along DPC1, the color-coded map “-2.4DPC1 vs 2.4DPC1” indicates that two monomers rotate in the same direction (black curved arrow) around a common axis through centers of monomers (black). Along DPC2, the color-coded map “-2.3DPC2 vs 2.3DPC2” indicates that two monomers rotate in opposite directions (black curved arrows) around two orthogonal axes (black). The rotation axis for the left monomer passes through the central stalk.

**Figure 5.**
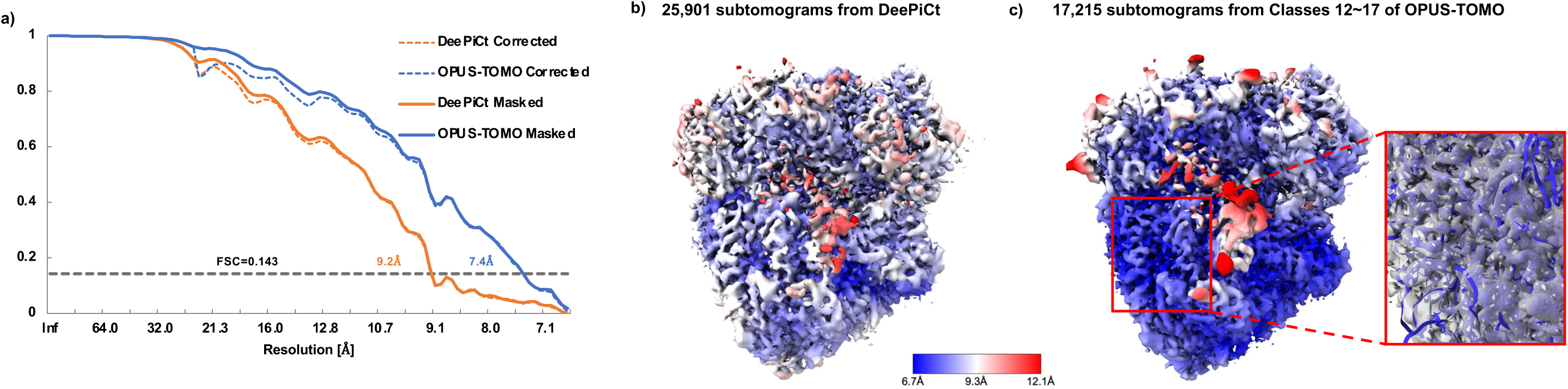
Reconstructions of the *S. pombe* 80S ribosome (data from EMPIAR-10988). **a**). Comparison between the gold standard FSCs of the subtomogram averages from different datasets. The name of FSC curve is a combination of prefix and suffix. The suffix represents the type FSC, where “Corrected” and “Masked”, represent corrected FSC and masked FSC, respectively. The prefix specifies the dataset for reconstructing the corresponding subtomogram average. “DeePiCt” represents the subtomograms annotated by expert from DeePiCt’s work (deposited in EMPIAR-10988). “OPUS-TOMO” represents the subtomograms from Classes 12∼17 classified by OPUS-TOMO from template matching result (**Extended Data Fig.**). **b)**. Subtomogram average of *S. Pombe* 80S ribosome using 25,901 subtomograms from DeePiCt’s annotations. Density map is contoured at 4𝜎 level, and colored by local resolutions. **c)**. Subtomogram average of *S. Pombe* 80S ribosome using 17,215 subtomograms from Classes 12∼17 by OPUS-TOMO. Density map is contoured at 4𝜎 level, and colored by local resolutions. A region (red box) where the subtomogram average from OPUS-TOMO presents higher local resolutions is shown in zoomed view, highlighting the fitting between the density and the secondary structure of *S. c* 80S ribosome (PDB code: 9BDP).

OPUS-TOMO possesses the unique capability to reveal the inter-subunit dynamics through its dynamics decoder. Using Class 16 (state 2) of ATP synthase dimer as the template (**Fig. 3a**), OPUS-TOMO reveals two orthogonal normal modes by traversing the first and second principal component of dynamics latent space (DPC1 and DPC2) and reconstructing traversed latent codes using dynamics decoder (**Fig. 4h**). The atomic structure for F_1_ Head and central stalk (PDB code: 6RDE) and the atomic structure for peripheral stalk (PDB code: 6RD4) was fitted into the density maps reconstructed by OPUS-TOMO along DPC1 and DPC2 (**Extended Data Fig. 4b∼c**). All the subunits in each monomer were fitted separately. Along DPC1, two monomers rotate in the same direction (**Extended Data Fig. 3b, Supplementary Video 2**), with the color-coded map for displacements between −2.4DPC1 and 2.4DPC1 indicating a common rotation axis passing through the centers of monomers (**Fig. 4h, Left**). In contrast, along DPC2, two monomers rotate in opposite directions (**Extended Data Fig. 3c, Supplementary Video 3**). The color-coded map for displacements between −2.3DPC2 and 2.3DPC2 indicates that the rotation axis for the left monomer passes through the central stalk, and the rotation axis for the right monomer passes through the peripheral stalk to the F_1_ Head.

### High-resolution Reconstruction of 80S Ribosome from Tomograms of Wild-type *Schizosaccharomyces* (*S.*) *pombe*

Another dataset for testing the performance of OPUS-TOMO consists of ten tomograms of wild-type *S. pombe* from DeePiCt’s work^39^ (EMPIAR-10988), where a set of 25,901 subtomograms for 80S ribosome was manually annotated by human expert (**Extended Data Fig. 4a**). This expert-annotated set from DeepPiCt’s work^39^ yielded a subtomogram average with a resolution of 9.2 Å after multiple rounds of 2D image warp, CTF and particle pose refinement in M^4^ (**Fig. 3a**) with low local resolutions (**Fig. 3b**).

Using template matching in pyTOM^16^ with a published *S. cerevisiae* 80S ribosome map^9^ (EMDB accession code: EMD-3228) as the template, we picked a total of 50,000 80S ribosomes (**Extended Data Fig. 4a**). We then performed structural heterogeneity analysis by training OPUS-TOMO with subtomograms and their orientations from the template matching result directly. The UMAP visualization of the 12-dimensional composition latent space learned by OPUS-TOMO shows distinct clusters, which we clustered into 20 classes by KMeans algorithm, and reconstructed 3D structure for each class by supplying cluster centers into the trained composition decoder of OPUS-TOMO (**Extended Data Fig. 4b∼c**). Classes 12∼17 on the left-hand side of UMAP visualization, totaling 17,215 subtomograms, show complete densities for 80S ribosomes, while the classes on the right-hand side of UMAP show densities for other species (**Extended Data Fig. 4b∼c**). All 17,215 subtomograms from Classes 12∼17 were imported into M^4^ for refinement. Under the same refinement settings for DeePict’s set, we obtained a subtomogram average of a resolution of 7.4 Å (**Fig. 3a**), with local resolutions approaching the Nyquist limit (6.7 Å) in most regions at the same contour level (4𝜎) (**Fig. 3c**). Using the atomic structure of *Saccharomyces cerevisiae* (*S. c*) 80S ribosome (PDB code: 9BDP)^40^ as a reference, the subtomogram average of Classes 12∼17 shows clear densities for the secondary elements in the atomic structure (**Fig. 3c**).

We also compared the subtomograms from template matching and Classes 12∼17 from OPUS-TOMO to DeePiCt’s annotations. Among the total 50,000 subtomograms from our template matching results, there were 21,351 subtomograms also found in DeePiCt’s annotations, of which 14,039 subtomograms were found in Classes 12∼17 of our OPUS-TOMO analysis (**Extended Data Fig. 4a and Extended Data Table 1, Left**). Thus, the overlapped subtomograms of our template matching results accounted for 43% (21,351/50,000) of DeePiCt’s annotations, while the overlap of Classes 12∼17 from OPUS-TOMO’s analysis with all subtomograms in Classes 12∼17 was improved to 82% (14,039/17,215). The distribution of these subtomograms in each tilt series is shown in **Extended Data Fig. 4d∼e** and **Extended Data Table 1, Left**. Clearly, OPUS-TOMO improves the percentage of ribosomes in the cluster over the original template matching result by a large degree.

In summary, a single-shot heterogeneity analysis by OPUS-TOMO on template matching result produced a significantly purified subtomogram set, of which 82% subtomograms overlap with human expert annotations, and improved the final resolution of 80S ribosomes by 1.8 Å, even with local resolutions approaching the Nyquist limit (6.7 Å) in most regions.

### Translocation-intermediate States and Dynamics of *S. pombe* 80S Ribosome

80S ribosome synthesizes proteins in eukaryotic cell by translating messenger RNA (mRNA) via a series of large-scale conformation changes facilitated by the elongation factors (eEFs). Specifically, it has three tRNA binding sites, A, P and E sites, spanning from 40S Head to 60S. In the elongation cycle, these sites are sequentially occupied and released by tRNAs and different eEFs, translating mRNA to peptide chains. We continue to reveal the dynamics and compositional heterogeneity which are crucial for understanding the protein synthesis *in situ* by training OPUS-TOMO on the M-refined subtomogram set purified by OPUS-TOMO. During training, the dynamics decoder leveraged a two-subunit model to capture the relative displacements between 40S and 60S subunits, in which each subunit is centered at its COM.

Clustering the composition latent space learned by OPUS-TOMO using KMeans^41^ results in 30 well-separated clusters (**Extended Data Fig. 5a**). The 3D structures for these classes are reconstructed by supplying the classes’ centroids into the trained composition decoder (**Extended Data Fig. 5b**). Within these 30 classes, Classes 26∼29 shows non-ribosome densities, Class 24 presents only the 60S subunit, remaining classes show complete densities for 80S ribosome. By close examination of the reconstructed classes, additional elongation translation cofactors bound to 80S ribosomes can be identified (**Fig. 6a**). For instance, it is easy to identify classes with eEF3 which is of ear shape and binds with 40S Head subunit (**Fig. 6a**). The density of eEF3 in Class 25 achieves a correlation coefficient (CC) of 0.75 by rigid body fitting of the atomic structure of eEF3 from an 80S ribosome structure (PDB code: 7B7D^42^) (**Fig. 6b, Right**). Two other eEFs for yeast revealed by OPUS-TOMO are eEF2 and eEF1A-tRNA which bear similar shapes and bind to 60S subunit at the same location, thus requiring reconstructions with high resolutions to distinguish. The density in Class 17 bound between 40S and 60S subunit is mapped to eEF1A-tRNA as it shows a triangular head (eEF1A) and smooth tail (tRNA), which achieves a CC of 0.74 with the structure of eEF1A bound with tRNA from an 80S ribosome structure (PDB code: 9BDP^40^) (**Fig. 6b, Left**). In contrast, the density in Class 25 at the same location is mapped to eEF2 as it shows a bigger head and rougher tail, which achieves a CC of 0.70 with the structure of eEF2 from an 80S ribosome structure (PDB code: 7OSM^33^) (**Fig. 6b, Middle**). Classes with eEF2 are found to be Class 18, 19, 21, 22 and 25, forming a big cluster in the UMAP visualization of composition latent space (**Extended Data Fig. 2a**). More impressively, different classes with eEF2 present a set of conformational changes in different subunits that clearly illustrate the role of eEF2 during the elongation translation cycle (**Fig. 6c∼d**). Comparing the density maps of Class 18 to Class 25 reveals a clockwise rotation of the 40S Head subunit and the eEF3 when transiting from Class 18 to Class 25 (**Fig. 6c, Middle**). The clockwise rotation of eEF3 leads to the binding of eEF3 with 60S subunit in Class 25 (**Fig. 6c, Left**). Meanwhile, the L1 stalk moves outward (**Fig. 6c, Middle**). The eEF2 subunit shifts rightward by a small amount (**Fig. 6c, Right**). By inspecting the densities of Class 18 near eEF2 and the A, P, E sites, it is identified as an translocation state with two tRNAs trapped in hybrid ap/P and pe/E transitory positions^33^ (**Fig. 6d, Top**). The identity of Class 18 is supported by rigid-body fitting of the atomic structure of the corresponding state (PDB code: 7OSM) and zoning near the eEF2 and the ap/P- and pe/E-site tRNAs (**Fig. 6d, Top**). In contrast, Class 25 is identified as a translocation state with two tRNAs trapped in classical P, E sites, which is supported by rigid-body fitting of the atomic structure of the corresponding state (PDB code: 9BDP) and zoning near the eEF2 and the P- and E- site tRNAs (**Fig. 6d, Middle**). The superposition of fitted atomic structures at selected zone shows a small rightward shift for two tRNAs during their transition from hybrid ap/P and pe/E transitory positions to classical P and E sites (**Fig. 6d, Bottom**). Therefore, the movements of eEF2 with neighboring tRNAs and 40S Head are shown to be in opposite directions, confirming the function of eEF2 as a ‘locking pawl’ that prevents the tRNAs in transitory positions rotating back with the 40S Head during translocation^33^ (**Fig. 6d, Bottom**).

**Figure 6.**
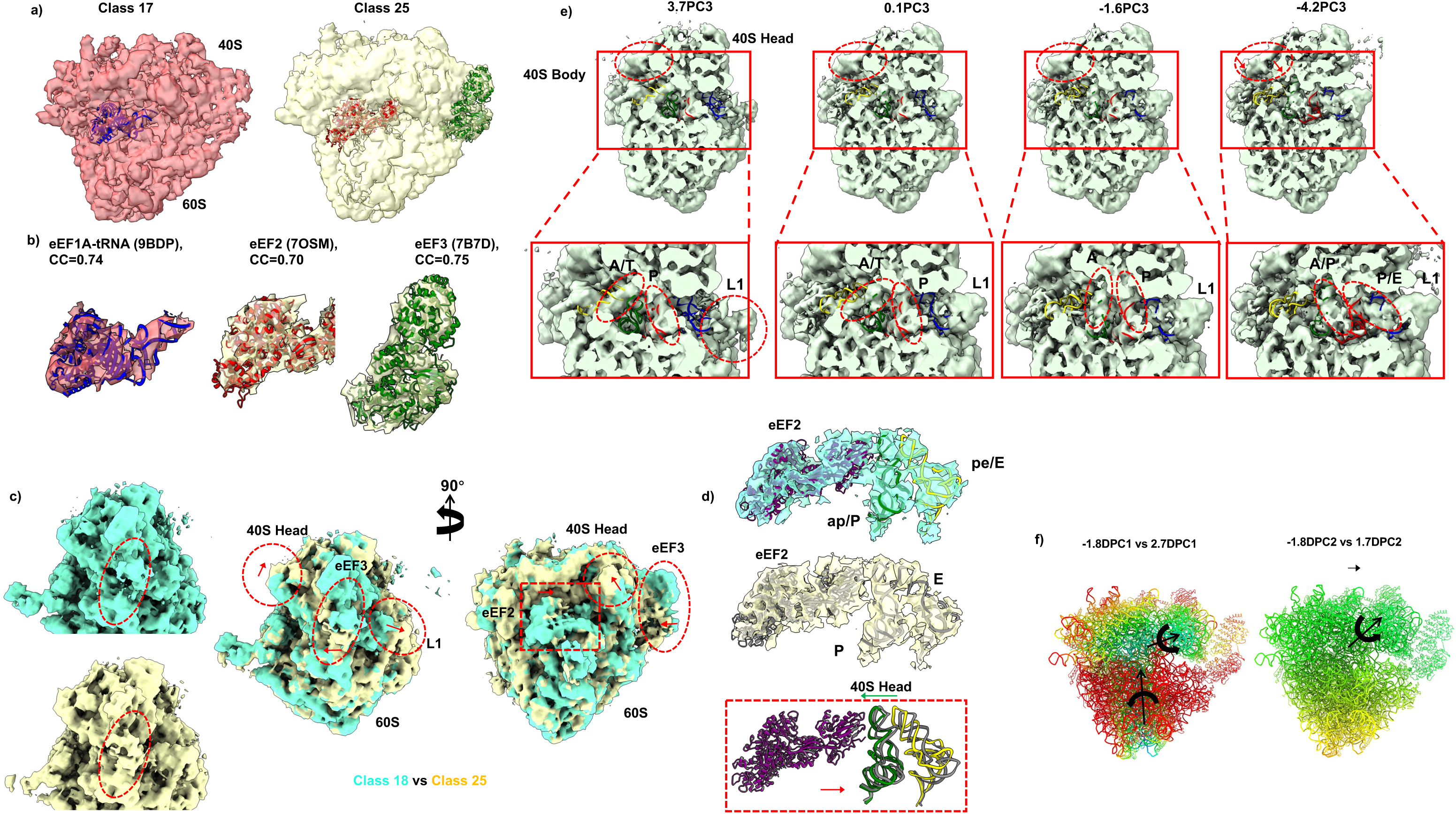
Translocation-intermediate states of the *S. pombe* 80S ribosome *in situ*. **a)**. 80S ribosomes with different elongation factors reconstructed by OPUS-TOMO at different class centroids (**Extended Data Fig. 4a**). Class 17 (transparent red) shows the structure of 80S ribosome with eEF1A-tRNA (blue) and eEF3, Class 25 (transparent yellow) shows the 80S ribosome with eEF2 (red) and eEF3 (green). **b)**. Densities of elongation factors from **a)** overlaid with their fitted structures (PDB code in parentheses). CC represents the correlation coefficient between densities and atomic models. **c)**. Conformational differences between translocation-intermediate states of *S. pombe* 80S ribosome with eEF2 and eEF3. (Left) The eEF3 in Class 18 (red dashed ellipse) is detached from 60S subunit, while the eEF3 in Class 25 (red dashed ellipse) is bound with 60S subunit. (Middle) Superposition of density maps reveals that 40S Head and the eEF3 bound to it rotate clockwise (red dashed ellipses and arrows) when Class 18 (cyan) transits to Class 25 (yellow). Meanwhile, the L1 stalk moves outward to open the E-site. (Right) Superposition of density maps further reveals that eEF2 moves rightward during the transition (red dashed rectangle and arrow). **d)**. Densities of eEF2 and two tRNAs from translocation-intermediate states in **c**) overlaid with the corresponding atomic structures. (Top) Densities from Class 18 overlay with the atomic structure of eEF2 and two tRNAs trapped in hybrid ap/P and pe/E sites (colored, PDB code: 7OSM). (Middle) Densities from Class 19 overlay with the atomic structure of eEF2 and two tRNAs trapped in P and E sites (grey, PDB code: 9BDP). (Bottom) Superposition of the structure fitted to Class 18 (colored) and the structure fitted to Class 19 (grey) reveals eEF2 acting as a pawl anchoring when two tRNAs transit from hybrid ap/P and pe/E sites to P and E sites (red arrow). Meanwhile, the 40S Head moves in reverse direction (green arrow) in relative to tRNAs. **e)**. Translocation-intermediate states revealed by progressing in the direction of PC3 in OPUS-TOMO’s composition latent space. The location of latent code where the density map is reconstructed by the composition decoder of OPUS-TOMO is specified as xPC3. The structures of tRNAs at A/T (blue, PDB code: 9BDP), A (green, PDB code: 7B7D), P (red, PDB code: 9BDP), and E (blue, PDB code: 9BDP) sites are shown as reference. Traversing PC3 shows the continuum of tRNA translocation (red dashed ellipses in zoomed views) to different sites (black texts). The red dashed circles in the top show 40S Head rotate counterclockwise (red arrow) in relative to 40S Body as indicated by the area of shadow. **f)**. Color-coded displacement maps reveal two distinct inter-subunit movements along different DPCs of OPUS-TOMO’s dynamics latent space. Residues in the structures are colored by their displacement distances along the corresponding DPCs (**Extended Data Fig. 5**), using rainbow palette in ChimeraX^57^ with a scale from 0 to 6.5 Å, where red indicates larger displacements and blue suggests smaller displacements. Along DPC1, the color-coded map “-1.8DPC1 vs 2.7DPC1” indicates that two subunits rotate in opposite directions (black curved arrows) around two orthogonal axes (black arrow). Along DPC2, the color-coded map “-1.8DPC2 vs 1.7DPC2” indicates that two subunits translate (black arrow) and rotate in the same direction (black curved arrow) around a common axis (black arrow).

The elongation cycle of 80S ribosome initiates with the delivery of tRNAs to a vacant A-site by eEF1A, transitions through various configurations including state with hybrid A/P- and P/E-site tRNAs, state with hybrid ap/P- and pe/E- site tRNAs which was revealed before, and state with P- and E-site tRNAs, ends with E-site tRNA exited^33^. The PCs of the composition latent space learned by OPUS-TOMO correspond to parts of the elongation cycle for 80S ribosome. For example, the positive end of PC3 (3.7PC3) shows a state with P-site tRNA and A/T tRNA bound to eEF1A cofactor, while the E site is opened by L1 stalk (**Fig. 6e**). Moving forward, the middle point along PC3 (0.1PC3) shows an intermediate state with A site half occupied, P site occupied and E site closed by L1 stalk (**Fig. 6e**). The next point along PC3 (−1.6PC3) shows a state with A and P sites occupied and E site closed by L1 stalk (**Fig. 6e**). The negative end of PC3 shows a state with hybrid A/P-site tRNA and P/E- site tRNAs binding with L1 stalk at E site, indicating the function of L1 stalk for facilitating the transition to this hybrid state (**Fig. 6e**). During the transition, the 40S Head and Body are rotating counter-clockwise as evidenced by the increasing gap between the 40S subunit and the top bar of red rectangles, which brings the A and P site in 40S Head close to the P and E site in 60S (**Fig. 6e, Top**). The transition along PC3 clearly demonstrates the translocation dynamics of tRNAs inside the 80S ribosome coupling with the counter-clockwise rotation of 40S (**Supplementary Video 4**).

For the 80S ribosome, using Class 17 as the template (**Extended Data Fig. 5a**), OPUS-TOMO reveals two distinct normal modes by traversing the first and second principal component of dynamics latent space (DPC1 and DPC2) and reconstructing traversed latent codes using dynamics decoder. The atomic structure for 80S ribosome (PDB code: 7B7D) was fitted into the density maps reconstructed by OPUS-TOMO along DPC1 and DPC2 (**Extended Data Fig. 6a∼b**). All the subunits were fitted separately. Along DPC1, the 40S subunit and 60S subunit rotate in the opposite directions (**Extended Data Fig. 6a**), leading to the twist of 80S ribosome (**Supplementary Video 5**). The color-coded map for displacements from −1.8DPC1 to 2.7DPC1 indicates that two subunits adopt two orthogonal rotation axes. 40S rotates around the axis passing through the 40S Body to 40S Head, while 60S rotates around the axis passing through the central axis of 60S and exhibits rotation in larger magnitude (**Fig. 6f, Left**). In contrast, DPC2 represents a degenerate movement for two subunits, in which they exhibit global translation and rotate around a common axis centering at the 40S subunit (**Extended Data Fig. 6b, Fig. 6f, Right, Supplementary Video 6**), resulting in a uniform gradient of displacement from 40S subunit to 60S subunit.

### Reconstruction of Fatty Acid Synthase from Tomograms of Wild-type *S. pombe*

To demonstrate the power of OPUS-TOMO under low data settings, we applied OPUS-TOMO to reconstruct a sparse populated species, fatty acid synthase (FAS) in tomograms of wild-type *S. pombe* (EMPIAR-10988)^39^. A total of 4,800 particles for FAS were picked using template matching method in pyTOM^16^ with a published *S. cerevisiae* FAS map^43^ (EMDB accession code: EMD-1623) as the template. We then performed 3D refinement for picked subtomograms using the low-pass filtered *S.cerevisiae* FAS map as the starting model, and obtained a low-resolution and not-so-continuous subtomogram average by RELION 3.0.8^9^. Given the 𝐷_3_ symmetry inherent to the FAS structure, we applied symmetry expansion to the refined pose parameters of subtomograms using RELION 3.0.8 as a preparatory step for structural heterogeneity analysis. Following this symmetry expansion, OPUS-TOMO was trained on subtomograms with the assumption that their projection angles correspond to one of the six equivalent angles satisfying 𝐷_3_ symmetry.

The UMAP visualization of the 10-dimensional composition latent space learned by OPUS-TOMO shows distinct clusters, which we clustered into 20 classes by KMeans algorithm, and reconstructed 3D structure for each class by supplying cluster centers into the trained composition decoder of OPUS-TOMO (**Extended Data Fig. 7a∼b**). Only Class 19 shows complete densities for FAS, and contains 221 particles.

The map of Class 19 displays higher-resolution features, for instance in regions marked by red solid and dashed box (**Fig. 7a, Left panel**), compared to the subtomogram average obtained by DeePiCt’s annotation^39^ (EMDB code: EMD-14422) (**Fig. 7a, Right panel**), when both maps are contoured at the same level. This is further supported by fitting the atomic structure of FAS (PDB code: 2VKZ^44^) into density maps (**Fig. 7b**). Judging by Fourier Shell Correlations (FSC)=0.143 between the atomic model 2VKZ and the density maps, Class 19 reconstructed by OPUS-TOMO reaches a resolution of 20.0 A#, while the subtomogram average in EMD-14422 from DeePiCt has a resolution of 27.4 A# (**Fig. 7c**). We also imported the subtomogram sets from both the expert-annotated set from DeepPiCt’s work^39^ and the Class 19 from OPUS-TOMO into M for further refinements while enforcing 𝐷_3_symmetry. The expert-annotated set from DeePiCt yielded a subtomogram average with a resolution of 18.5 Å after multiple rounds of 2D image warp, CTF and particle pose refinement in M^4^ (**Fig. 7f**) with low local resolutions (**Fig. 7d**). The refinement protocol on the Class 19 from OPUS-TOMO yielded a subtomogram average with a resolution of 14.0 Å (**Fig. 7f**) with higher local resolutions (**Fig. 7e**).

**Figure 7.**
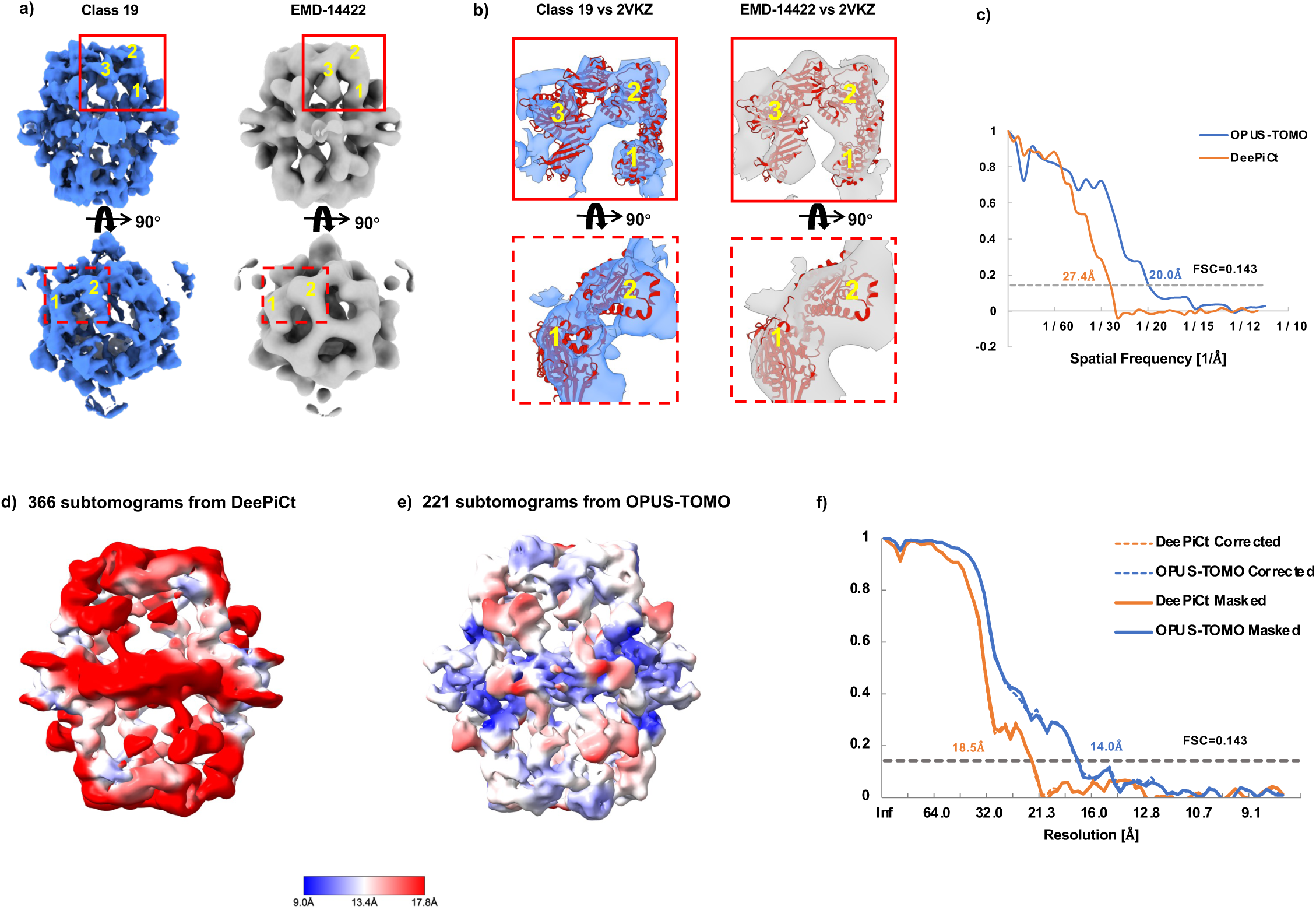
Reconstructions of the *S. pombe* FAS (data from EMPIAR-10988). **a)**. Comparison between the FAS reconstructed by OPUS-TOMO (Class 19) and the deposited density maps for FAS (EMD-14422). Side and top views of these EM maps are presented. Class 19 reconstructed by the composition decoder of OPUS-TOMO shows complete densities for FAS, and EMD-14422 shows the subtomogram average of subtomograms picked by DeePiCt^39^. Class 19 presents structural features at a higher resolution than EMD-14422 (red boxes where separate domains are labeled by texts). **b).** Zoomed views of the boxed regions in **a)** with the atomic structure of FAS with PDB accession code 2VKZ^44^ fitted into density maps. Separate domains are labeled by texts as in **a)**. **c**) Model-map FSCs between the atomic structure of FAS (PDB code: 2VKZ) and the density maps from OPUS-TOMO and DeePiCt. **d)**. Subtomogram average of *S. pombe* FAS using 366 subtomograms from DeePiCt’s annotations. Density map is contoured at 5𝜎 level, and colored by local resolutions. **e)**. Subtomogram average of *S. pombe* FAS using 221 subtomograms from Class 19 by OPUS-TOMO. Density map is contoured at 5𝜎 level, and colored by local resolutions. **f**). Comparison between the gold standard FSCs of the subtomogram averages of DeePiCt’s annotations and OPUS-TOMO. The FSC curves are named using the same convention as in **Fig. 5a**.

Given the sparsity of FAS population, picking FAS from the template matching results is similar to finding a needle in a haystack. Among the total number of 4,800 subtomograms from our template matching results, there were 115 subtomograms that overlapped with DeePiCt’s expert annotation result, among which 89 subtomograms were found in Class 19 of our OPUS-TOMO analysis (**Extended Data Table 1, Right; Extended Data Fig. 7c**). The distribution of these subtomograms in each tilt series is shown in **Extended Data Fig. 7d** and **Extended Data Table 1, Right**. **Extended Data Fig. 7e** shows the significantly enhanced ratios of subtomograms overlapping with DeePiCt’s expert annotation result for Class 19 from OPUS-TOMO for each tilt series than for the original template matching result. Furthermore, although the portion of subtomograms that were included in both OPUS-TOMO and DeePiCt was relatively small, both methods faithfully re-captured the essential features of FAS. This is exemplified by the similarity of subtomograms from tilt series TS_045, where the 16 subtomograms in Class 19 of our analysis and the 18 subtomograms in DeePiCt’s expert annotation result, in spite of no overlaps, exhibited similar densities corresponding to the FAS complex (**Extended Data Fig. 7f**).

### High-resolution Reconstruction of 70S Ribosome from Tomograms of *Mycoplasma (M.) Pneumoniae*

To further test the capability of OPUS-TOMO for facilitating high-resolution reconstruction, we applied OPUS-TOMO to the reconstruction of 70S ribosomes from *M. Pneumoniae* tomograms^45^ (EMPIAR accession code: 10499). The tilt series were automatically aligned and reconstructed as tomograms using AreTOMO^46^. Using the *M. Pneumoniae* 70S ribosome structure^45^ (EMD accession code: EMD-11999) which was lowpass filtered to 40 Å as a template, for each of the 64 tomograms, the 750 top-scored subtomograms in the 8x binned tomograms were picked by template matching in pyTOM^16^. The coordinates of picked subtomograms and tilt alignment parameters were imported into WARP^6^ to reconstruct subtomograms and estimate their 3DCTF parameters. The reconstructed subtomograms were aligned by pyTOM^16^, and the aligned coordinates and poses were imported into M^4^ for further refinement. The raw template matching set converged to a resolution of 4.8 Å after multiple rounds of 2D image warp, CTF and particle pose refinement (**Fig. 8a, 8c**).

**Figure 8.**
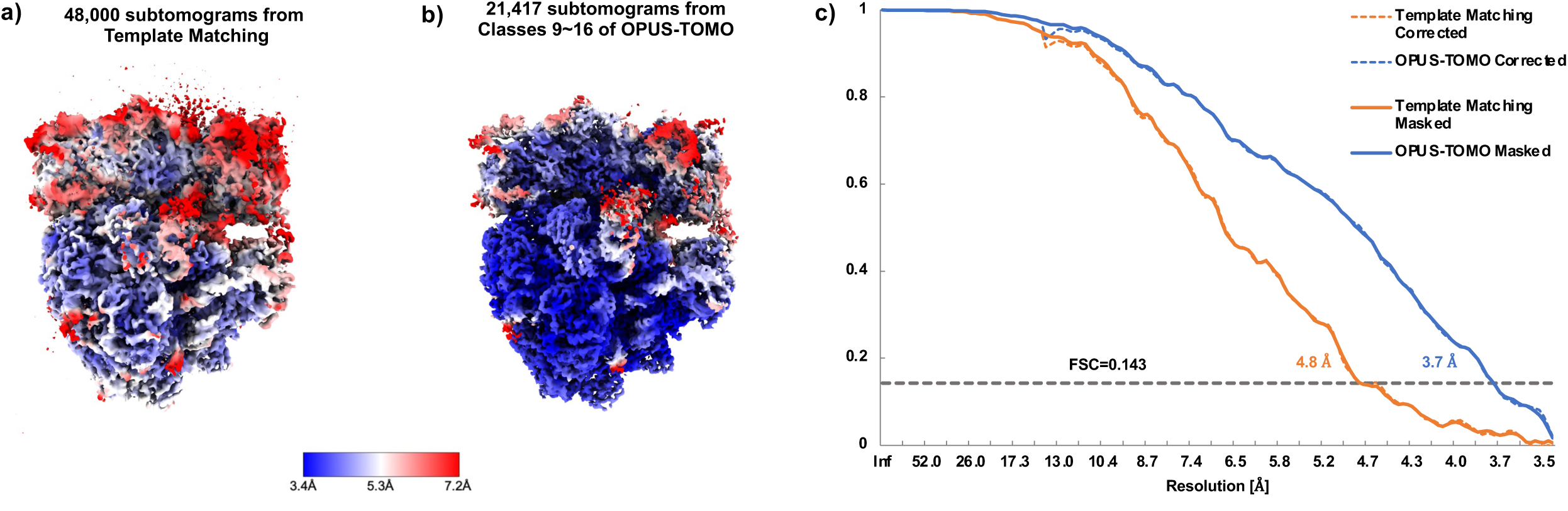
Reconstructions of the *M. pneumoniae* 70S Ribosome (data from EMPIAR-11843). **a)**. Subtomogram average of *M. pneumoniae* 70S ribosome using 48,000 subtomograms from template matching. Map is contoured at 4𝜎 level, and colored by local resolutions. **b)**. Subtomogram average of *M. pneumoniae* 70S ribosome using 21,417 subtomograms from Classes 9∼16 by OPUS-TOMO. Map is contoured at 4𝜎 level, and colored by local resolutions. **c**). Comparison between the gold standard FSCs of the subtomogram averages of template matching and OPUS-TOMO. “Template Matching Corrected” represents the corrected FSC of the subtomogram averages of the original template matching set. “OPUS-TOMO corrected” represents the corrected FSC of the subtomogram averages of Classes 9∼16 given by OPUS-TOMO. “Template Matching Masked” represents the masked FSC of the subtomogram averages of original template matching set. “OPUS-TOMO Masked” represents the masked FSC of the subtomogram averages of Classes 9∼16 given by OPUS-TOMO.

The refined coordinates were imported into WARP to reconstruct subtomograms, which together with their orientations and CTF parameters determined by M were then supplied into OPUS-TOMO for heterogeneity analysis. The UMAP^47^ visualization of the 12-dimensional composition latent space learned by OPUS-TOMO shows distinct clusters, which we clustered into 30 classes by KMeans algorithm, and reconstructed 3D structure for each class by supplying cluster centers into the trained composition decoder of OPUS-TOMO (**Extended Data Fig. 8**).

Classes 9∼16 in UMAP visualization show strong and continuous densities for 70S ribosomes, Classes 19∼20 show weak and noisy densities for 70S ribosomes, while the remaining classes in UMAP don’t resemble 70S ribosome. Combining Classes 9∼16 yielded 21,417 subtomograms in total, which represent 45% (21,417/48,000) of the subtomograms picked by template matching. The subtomograms of Classes 9∼16 were imported into M^4^ for refinement, and converged to a resolution of 3.7 Å after multiple rounds of 2D image warp, CTF and particle pose refinement (**Fig. 8c**), with local resolutions approaching the Nyquist limit (3.4 Å) in most regions at the same contour level (4𝜎) (**Fig. 8b**).

### Translocation-intermediate States of *M. Pneumoniae* 70S Ribosome

For comparison to existing approaches on structural heterogeneity analysis using high-resolution STA result, we tested OPUS-TOMO in analyzing 70S ribosomes from *Mycoplasma (M.) Pneumoniae* ^30,45^ (EMPIAR accession code: 11843) at a resolution of ∼3.5 Å, which is the same dataset used in the analysis by tomoDRGN. The subtomograms of size 210^3^ were reconstructed from the M-refined particle tilt series by RELION 3.0.8^48^, and were then supplied into OPUS-TOMO for heterogeneity analysis using their orientations and CTF parameters determined by M. For comparison, the dynamics decoder only models the rotation and translation of the whole complex during training.

The UMAP^47^ visualization of the 12-dimensional composition latent space learned by OPUS-TOMO shows distinct clusters, which we clustered into 30 classes using KMeans algorithm, and reconstructed 3D structure for each class by supplying cluster centers into the trained composition decoder of OPUS-TOMO (**Extended Data Fig. 9a∼b**). Significant conformational changes of 70S ribosome can be revealed by comparing different classes. For example, large joint conformational changes of the head of 30 subunit and L1 stalk were identified by comparing the density maps of Class 16 and Class 19 (**Fig. 9a**). Class 11 and Class 23 show densities for the EF-Tu cofactor between the body of 30S subunit and L10 stalk (**Extended Data Fig. 9b**). By comparing Class 11 to a state without the EF-Tu cofactor, Class 19, we can observe that the binding of EF-Tu cofactor induces large displacements of RNAs in the body of 30S subunit (**Fig. 9b**). Besides, we can also observe densities for L7/L12 NTD dimer of dimers in Class 11 and Class 19, which are consistent with previous study by tomoDGRN^30^ (**Fig. 9b, red solid circle**).

**Figure 9.**
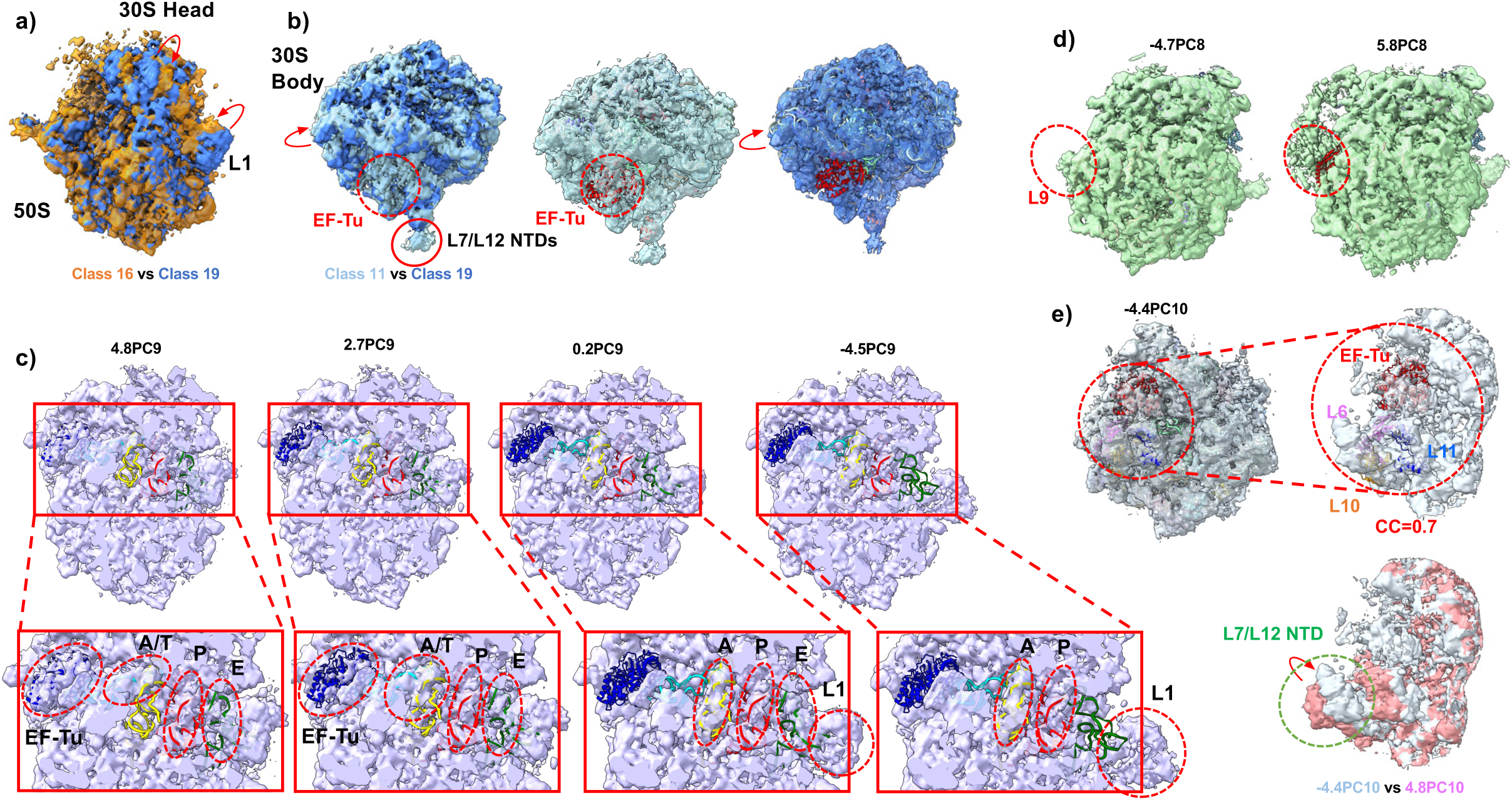
Translocation-intermediate states of the *M. pneumoniae* 70S ribosome *in situ*. **a)**. Superposition of density maps for different translation-intermediate states revealing the large-scale displacement between the 30S Head and L1 stalk (red arrow) when transiting from Class 19 (blue) to Class 16 (brown). **b)**. (Left) Superposition of density maps for different translocation-intermediate states revealing the conformational change of the 30S Body (red arrow) induced by the binding of EF-Tu cofactor (red dashed circle) in Class 11 (light blue). (Middle) The density map of Class 19 overlaid with the structure of 70S ribosome (colored, PDB code: 7PHA^49^) with EF-Tu (red). (Right) The density map of Class 11 overlaid with the structure of 70S ribosome with EF-Tu to show the absence of densities of EF-Tu. Red circle marks the densities for L7/L12 NTDs. **c)**. Translocation-intermediate states revealed by the progressing in the direction of PC9 in OPUS-TOMO’s composition latent space. The locations of latent codes are specified as xPC9. The structures of A- (yellow), P- (red) and E-site (green) tRNAs from *T. thermophilus* Cm-bound 70S ribosome (PDB code: 6ND5^58^) and the structures of EF-Tu (blue) and A/T-site tRNA (cyan) from 70S ribosome in Cm-treated *M. pneumoniae* cell (PDB code: 7PHA^49^) are shown as reference. Traversing PC9 shows the continuum of tRNA translocation (red dashed ellipses in zoomed views) to different sites (black texts). **d)** Occupancy change of L9 subunit identified by OPUS-TOMO using latent codes at different locations along PC8. The locations of latent codes are specified as xPC8. The density map reconstructed at −4.7PC8 overlaid with the structure of 70S ribosome (colored, PDB accession code: 7PHA^49^) shows densities for L9 (red dashed ellipse). The density map reconstructed at 5.8PC8 shows no densities for L9 (red). **e)** Occupancy change of EF-Tu cofactor and the associated conformational changes identified by OPUS-TOMO using latent codes at different locations along PC10. The locations of latent codes are specified as xPC10. The atomic structure of 70S ribosome with EF-Tu cofactor (PDB accession code: 7PHA^49^) is fitted into density map at −4.4PC10. In the zoomed view (red dashed circle), the densities are in close agreement with the structures of EF-Tu (red), L6 (purple), L11 (blue) and L10 (brown). The CC between the atomic structure of EF-Tu and the corresponding densities from density map at − 4.4PC10 reaches 0.7. The superposition of density maps at −4.4PC10 and 4.8PC10 reveals the displacement of the L7/L12 NTD (red arrows, green dashed ellipse).

Principal component analysis in the 12-dimensional composition latent space learned by OPUS-TOMO automatically reveals a set of spatially correlated structural changes. Traversing along the PC9 reveals a part of elongation cycle: the cycle starts from a state with EF-Tu-A/T-site tRNA and P- and E-site tRNA (4.8PC9), through an intermediate state with partly occupied A-site and P- and E-site tRNAs (0.2PC9), transiting to a state with A-, P- and E-site tRNAs, culminating in a state with only A- and P-site tRNAs (−4.5PC9) (**Fig. 9c**). Specifically, PC9 captures the delivery of tRNA to A-site by EF-Tu cofactor and the release of E-site tRNA facilitated by L1 stalk (**Fig. 9c** and **Supplementary Video 7**). Additionally, traversing PC9 reveals the orchestration of different structural elements in 70S ribosome during the transition. The RNA above the EF-Tu cofactor in the body of 30S subunit is shifting downwards as the EF-Tu cofactor being released, and the head of 30S subunit is shifting rightwards as the L1-stalk opens the E-site (**Supplementary Video 7**). However, it is worth noting that the spatially correlated states in elongation cycle revealed by traversing PC9 are not necessarily causal.

Traversing along the PC8 reveals the occupancy change of L9 on 50S subunit (**Fig. 9d**). Using the atomic models of 70S ribosomes in Cm-treated *M. pneumoniae* cell^49^ (PDB accession code: 7PHA) as reference, the densities for L9 disassociate from the 50S subunit when transiting from −4.7PC8 to 5.8PC8, while significant amount of densities appear at the neighboring region of L9.

Traversing along the PC10 reveals occupancy change of EF-Tu cofactor and the conformational change for L7/L12 NTDs (**Fig. 9e**). The EF-Tu cofactor without tRNA presents in the density map reconstructed at −4.4PC10. The correlation between the atomic structure of EF-Tu cofactor and the density map at −4.4PC10 reaches 0.7. The atomic structures of neighboring subunits, L6, L10 and L11, also fit with the neighboring densities. When transiting to 4.8PC10, the EF-Tu cofactor disappears while the densities for L7/L12 NTDs flexing away from the region hosting EF-Tu cofactor (**Fig. 9e**).

Traversing along the PC1 reveals occupancy change of peptide at the peptide exit tunnel, which is located between L29 and L22, and supported by L24 (**Extended Data Fig. 9c**). In the density map reconstructed at −10.0PC1, a tail-like density for peptide chain grows out from the peptide exit tunnel. In contrast, in the density map reconstructed at −2.3PC1, the tail-like density disappears.

The structural heterogeneity of 70S ribosome in Cm-treated *M. pneumoniae* cell was previously studied by cryoDRGN-ET^31^ and tomoDRGN^30^ (**Supplementary Video 8**). In cryoDRGN-ET^31^, the translation dynamics is reconstructed by using the latent codes manually selected from latent space **(Supplementary Video 8, Left panel**). In tomoDRGN^30^, the translation dynamics is reconstructed by manually selecting translation states among 100 structures at KMeans centroids **(Supplementary Video 8, Right panel**). By contrast, OPUS-TOMO reconstructed the translation dynamics by traversing PC9 without any human selection, and achieves much higher resolution and continuity **(Supplementary Video 7)**. This comparison demonstrates that OPUS-TOMO excels in associating spatially correlated structural changes to the principal components of latent space.

We further compared the density maps of 70S ribosome with EF-Tu-tRNA in translational dynamics reconstructed by different methods (**Extended Data Fig. 9d**). The contour levels of density maps were normalized to show similar volumes of densities for EF-Tu-tRNA in all density maps. Among all methods, the density map produced by OPUS-TOMO exhibits the best overall map quality and stronger and more complete densities for EF-Tu-tRNA (**Extended Data Fig. 9d**).

Lastly, we quantitatively compared the performances of structural heterogeneity analysis of different methods w.r.t the number of training data. We selected the subtomograms for 70S ribosome from the first eight tomograms in *M. Pneumoniae* ^30,45^ dataset (EMPIAR accession code: 11843), which consists of 2,990 subtomograms. We trained tomoDRGN and OPUS-TOMO for structural heterogeneity analysis using the same data by 50 epochs. The latent spaces learned by both methods at the last epoch were clustered into twenty classes. Clearly, by UMAP visualization, the composition latent space learned by OPUS-TOMO shows well separated clusters (**Extended Data Fig. 10a**). In these classes, Classes 0-5 and Class 19 are clearly noise, and Class 18 is 50S subunit, while the remaining classes show complete densities for 70S ribosome (**Extended Data Fig. 10b**). We further validated the clustering results by reconstructing the subtomogram averages using subtomograms from Class 0, Class 18 and Class 13. For each class, the subtomogram averages reconstructed by *relion_reconstruct* is consistent with the density map reconstructed by the composition decoder of OPUS-TOMO (**Extended Data Fig. 10c**). In contrast, the latent space learned by tomoDRNG shows classes with less separation (**Extended Data Fig. 10d**), and the density maps for different classes reconstructed by the decoder of tomoDRGN failed to reveal the non-ribosomes and 50S subunit compared to OPUS-TOMO’s result (**Extended Data Fig. 10e**). Therefore, using only 2,990 subtomograms, OPUS-TOMO can still reliably separate non-ribosomes and 50S subunit from 70S ribosome. This comparison demonstrates the robustness of OPUS-TOMO for structural heterogeneity analysis under smaller datasets, which is critical for analyzing sparse populated species in cell, and for analysis with a small number of tomograms.

## Discussion

Structural heterogeneity presents a fundamental challenge in cryo-ET data processing, limiting both resolution and biological interpretation. Here we present OPUS-TOMO, a deep learning framework that facilitates the high-resolution cryo-ET structure determination and the investigation of *in situ* biological processes for biomolecules. By encoding subtomograms into two sperate continuous low-dimensional latent spaces, composition latent space and dynamics latent space, from which the per-particle 3D structures and conformations for subtomograms can be decoded, OPUS-TOMO enables multiscale analysis of structural heterogeneity while overcoming key limitations of conventional methods.

Compared to traditional 3D classification approaches, OPUS-TOMO offers several significant advantages. It produces subtomogram sets for target specie from template matching result in substantially reduced computational time. Further refinements on the subtomogram sets from OPUS-TOMO yield higher-resolution reconstructions compared to those annotated by human expert via multiple rounds of classifications, frequently reaching the Nyquist limit of dataset. Most importantly, OPUS-TOMO scales to reconstruct structural continua, providing a continuous view of *in situ* structural heterogeneity at the molecular level compared to traditional classification methods. This capability is particularly valuable for uncovering functionally important transient states^20^.

Compared to existing deep learning based structural heterogeneity methods, OPUS-TOMO presents unique capactities to resolve speice-level heterogeneity in template matching results and model the large-scale inter-subunit movement of biomolecules using its dynamics decoder based on low-dimensional physical model. OPUS-TOMO often reveals the inter-subunit movements as orthogonal modes along different DPCs of the dynamics latent space, resmebling the low-frequency large-scale deformational motions revealed by normal mode analysis^20^. Therefore, the ability of dynamics decoder to reveal normal mode motions strongly endorses the power of OPUS-TOMO to directly capture functionally important motions of macromolecules using cryo-ET data. The distillation of large-scale inter-subunit movements in cryo-ET data into dynamics latent space also faciliates the modelling of intra-subunit changes by the composition decoder of OPUS-TOMO, improving the accuracy of composition latent code. Besides, the structural heterogeneity analysis by OPUS-TOMO is robust even with a much smaller dataset than required by existing methods, such as tomoDRGN.

The effectiveness of OPUS-TOMO stems from its smooth latent spaces and accurate latent codes. By incorporating both KL divergence from 𝛽-VAE^28,36^ and structural disentanglement priors from OPUS-DSD^29^, we ensure the latent space maintains smooth, physically meaningful relationships between structural states. Hence, OPUS-TOMO can reconstruct structural continua by simple linear interpolation in the latent space, as demonstrated by our visualization of the ATP synthesis process by ATP synthase and tRNA translocation dynamics by 80S ribosome. By contrast, existing approaches, such as tomoDRGN^30^ and cryoDRGN-ET^31^, implement much more complex analysis pipelines with a much larger number of samples in latent space in order to uncover similar structural heterogeneities, such as MAVEn analysis in tomoDRGN^30^ and the ensemble analysis in cryoDRGN-ET^31^. The accruacy of the latent code is supported by sub-nanometer resolution reconstruction of ATP synthase dimer and different ribosomes using classes clustered by KMeans algorithm in composition latent space. In addition, OPUS-TOMO discovers functionally important intermediate states such as the 80S ribosome with hybrid ap/P- and pe/E-site tRNAs^33^ which was not revealed by existing deep learning based approaches. Comparing different neighbouring states in the composition latent space of OPUS-TOMO frequently reveals biological insights for functionally important subunit in the macromelcules.

The implications of this work extend beyond methodological advances. By providing accurate analysis of structural heterogeneity with reduced human intervention for cryo-ET datasets, OPUS-TOMO makes high-resolution cryo-ET more accessible. Its ability to resolve continuous structural transitions offers new opportunities to study dynamic processes *in situ* and discover unstable intermediate states that are impossible to obtain by conventional experiments. The reconstructed intermediate states also provide much-needed insights to the functional roles of subunits in macromolecules. As cryo-ET moves towards broader biological systems, tools like OPUS-TOMO that can efficiently parse structural heterogeneity will be essential for speeding up structure determination and deepening our understandings about biomolecules *in situ*.

## Methods

### Deformation field

OPUS-TOMO represents the 3D structure of biomolecule by explicitly modelling its composition and deformation. The composition of a 3D structure can be represented by a volume on a discrete 3D grid, which is referred as template, and the deformation in relative to the template can be represented by a 3D deformation field. Each voxel in the deformation field records its original position in the template volume. The 3D conformation in OPUS-TOMO is sampled from the template according to the deformation field. Formally, the 3D conformation can be given by the composition of two functions *V*(***x*** + 𝒈(***x***)): ℝ^3^ → ℝ, where *V*(·) is the template which maps a grid point ***x*** to a voxel value, and 𝒈(***x***): ℝ^3^ → ℝ^3^ is the deformation field which record the displacement between the voxel ***x*** in the deformed conformation and its source voxel ***x*** + 𝒈(***x***) in template. Next, we give a detailed definition about the deformation field.

In OPUS-TOMO, the deformation of macromolecule is composed of individual movements of multiple subunits. The 3D deformation field is constructed through several steps. Firstly, a 3D consensus model is segmented into a set of pre-defined subunits, each of which has a specific shape with masses inside and undergoes rigid-body movement. Mathematically, a subunit with center 𝒄_𝑖_ ∈ ℝ^3^ can be defined as a function on the 3D grid, which is of the form

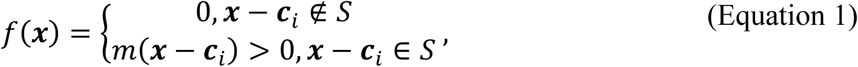

where 𝑆 refers to the set of grid points belongs to the subunit, and 𝑚(***x***) is the mass at point ***x***. We can further define some shape statistics about the subunit. For a subunit with center 𝒄_𝑖_ ∈ ℝ^3^, and suppose its principal axes form a matrix 𝑃_𝑖_, where each row is one principal axis, its semi-principal diameters can be defined as

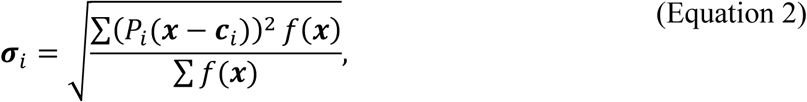

where the summations are over all grid points in the 3D grid. For a grid point ***x*** = {𝑥, 𝑦, *z*} ∈ ℝ^3^ in a subunit 𝑖, suppose the subunit undergoing a rigid-body displacement where the subunit is rotated by a rotation matrix 𝑅_𝑖_ ∈ 𝑆𝑂(3), and translated by the translation vector 𝒕_𝑖_ ∈ ℝ^3^, let the center of mass (COM) of the subunit be 𝒄_𝑖_, the rigid-body displacement of this point can be expressed as

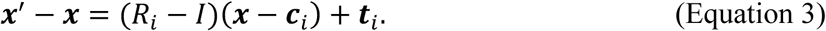

For a grid point in a large complex which is comprised of multiple subunits, each of which undergoes independent movement, its displacement should be the combination of those movements. We propose to blend the movement from a subunit at a grid point according to its distance w.r.t the center of subunit. Specifically, for a grid point ***x***, suppose the principal axes of subunit 𝑖 form a matrix 𝑃_𝑖_ as preceding, the displacement incurred by the movement of subunit 𝑖 decays as,

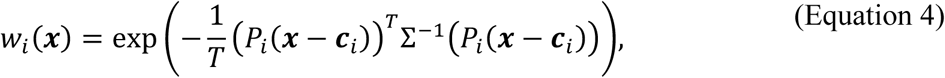

where 𝑥_j_is the 𝑗th component of the grid point ***x***, Σ is a diagonal matrix with its diagonal element Σ_jj_= 𝜎_𝑖,j_, *i.e.*, the semi-principal diameter of subunit 𝑖 along the 𝑗th axis, and 𝑇 is a constant for controlling the falloff of the weight function, which is set to be 5. The weight function should be normalized if there are *n* subunits in the complex. The normalized weight function can be expressed as,

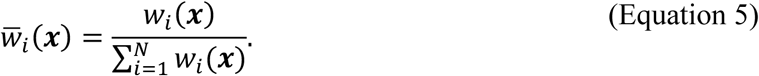

In a complex with *n* subunits, the displacement of a grid point is the linear combination of movements of different subunits, which can be written as,

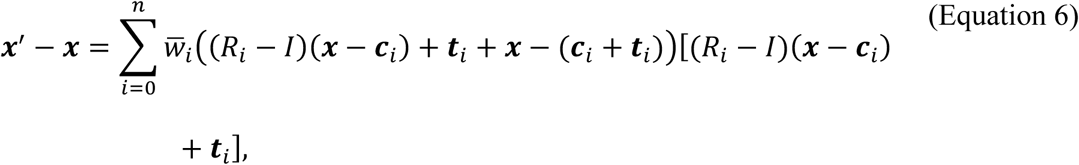

where *w̄*_𝑖_ controls the scale of the contribution of the movement of subunit 𝑖. We hence defined the deformation field 𝒈, with which the deformed grid can be represented as ***x***^′^ = ***x*** + 𝒈(***x***). In OPUS-TOMO, the rigid-body transformation parameters, 𝑅_𝑖_ and 𝒕_𝑖_, are predicted by the neural network. They should be restrained to rule out unrealistic deformations. We assume that the translational modes of subunits are generated by rotating COMs of subunits in relative to the COM of a common fixed subunit, i.e., 𝒕_𝑖_ = Y𝑅_𝑖,𝑡_ − 𝐼ZY𝒄_𝑖_ − 𝒄_𝑓_Z, where the COM 𝒄_𝑖_ is rotated by 𝑅_𝑖,𝑡_ centering at COM 𝒄_j_, and the translation of subunit can be obtained as the difference between the rotated COM and the original COM. The translation defined in this way retains the distance between COMs of subunits. The determination of the translation of each subunit comes down to estimating two rotation matrices. We can further limit the range of motion by restraining the rotation matrix to be an identity matrix, which inevitably hurts the expression power of the deformation model. To maximize the capability of the deformation model under the identity restraint, the rigid-body movement should be defined in an appropriate reference frame, *i.e.*, the *z* axis should align with the rotation axis. The rotation matrix which generates the translation of COM of subunit can be decomposed as 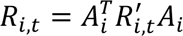, where the rotation matrix 𝐴_𝑖_ aligns the rotation axis of the COM of subunit to the *z* axis, and 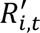 is the rotation matrix predicted by neural network. The rotation matrix of the subunit can be decomposed similarly as 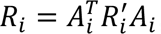, where 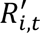 is the rotation matrix predicted by neural network. Our method adopts quaternion to represent rotation. The neural network in dynamics decoder outputs the last three dimensions of the quaternion, while the first dimension of quaternion is fixed to be a positive value, 16, which can regularize it to be an identity rotation.

### Subtomogram Formation Model

To reconstruct a 3D tomogram for the cellular sample, cryo-ET collects a series of 2D projections with different tilt angles. The 2D tilt series for a structure can be backprojected into a 3D volume to reconstruct a 3D subtomogram according their tilt angles. As in cryo-EM, the Fourier transform of image collected in cryo-ET is attenuated by the microscope CTF. We define a subtomogram formation model describing the effects of the CTF in 3D Fourier domain and the conformational changes in 3D structure.

For a subtomogram 𝑋_𝑖_, let the template volume be *V*, suppose 𝑋_𝑖_ is deformed by the deformation field 𝒈 and rotated from *V* by rotation matrix 𝑅_𝑖_ with Euler angle 𝜃, the Fourier transform of a subtomogram has the following relation w.r.t the Fourier transform of the template volume,

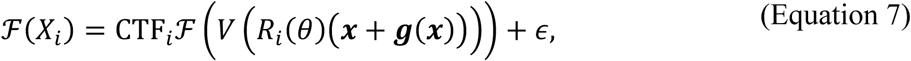

where ***x*** is the 3D coordinate of a voxel, CTF_𝑖_ is a 3D function in Fourier domain that describes the effects of CTF for the 2D images in the tilt series, ℱ and ℱ^−1^ denote Fourier transform and inverse Fourier transform, and 𝜖 is the 3D complex Gaussian noise, which is assumed to be independent and zero-mean for each lattice point. Specifically, CTF_𝑖_ is be obtained by backprojecting the 2D CTFs of all images in the tilt series into an empty 3D volume in Fourier domain. Each 2D CTF is placed as a central slice in the 3D CTF, whose orientation is determined by its tilt angle. Reconstructing the CTF model for subtomograms by backprojection naturally handles the missing wedges as the region which is not covered by tilt series remains empty, and preserves the tilt variations of CTFs by scattering CTF from different tilts to different voxels. Another characteristic of CTF in cryo-ET is the large variations in space due to the tilting and the thickness of specimen. The tilting of the specimen holder causes the specimen swing up (overfocus) or down (underfocus) on either side of the tilt axis^13^. In addition, particles in the specimen are distributed in different heights along 𝑍 axis. Hence, subtomograms extracted from different locations of the tomogram have different defocus. Various methods have been proposed to correct the CTF gradient in cryo-ET^13^. In this paper, we implemented a backprojection algorithm for reconstructing 3D CTF from the per-particle CTF parameters estimated by WARP^6^. Specifically, OPUS-TOMO reads the defocus parameters from Self-defining Text Archiving and Retrieval (STAR) format^50^ files and reconstructs the 3DCTF for each subtomogram *ad hoc* during training.

To capture the structural heterogeneity, the reference volume in (Equation 7) is defined on a per-particle basis. We assume that the structures of subtomograms reside in a low-dimensional latent space. Since a structure can be viewed as a combination of specific composition and conformation, the latent space is assumed to be composed of a low-dimensional vector 𝒛 describing its composition and a low-dimensional vector 𝒛^*d*^ describing its conformation. Given a neural network *V* that transforms 𝒛 into a 3D density map, and a neural network 𝒈 that transforms 𝒛^*d*^ into the deformation field, the Fourier transform of the subtomogram of the particle can be expressed as

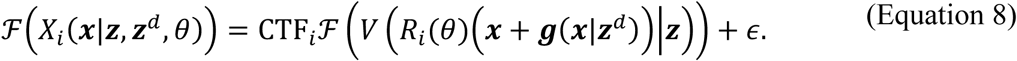

(Equation 8) defines the subtomogram formation model for the particle in cryo-ET which describes the effects of both spatial CTF variations and composition and conformation heterogeneity.

The CTF can be decomposed as a phase term, sign(CTF), which flips the sign of Fourier transform of subtomogram and leads to the spatial displacement of structure factor spacing in the image^51^, and an amplitude term, abs(CTF), which attenuates the frequency near zeros of the CTF. Hence, multiplying both sides of (Equation 8) by the phase of CTF, it can be rewritten as,

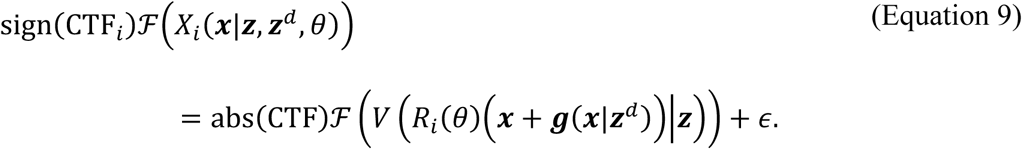

The resulting phase flipped subtomogram can be viewed as a focused subtomogram with reduced image spacing displacement, which is of higher signal to noise ratio and used for training OPUS-TOMO in this paper.

### Training Objective

The main objective of OPUS-TOMO is learning the per-particle structure for a subtomogram with pose determined by template matching or subtomogram averaging. The neural networks are trained by minimizing the difference between the subtomogram reconstructed from the 3D structure and the phase flipped experimental subtomogram 𝑋_𝑖_, that is,

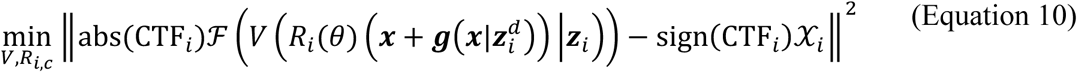

where 𝒳_𝑖_ is the Fourier transform of 𝑋_𝑖_, 𝒛_𝑖_ is the composition latent code for the template volume of subtomogram 𝑖, and 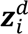 is the dynamics latent code for the deformation of subtomogram 𝑖. The training objective in (Equation 10) is not complete yet since the latent codes 𝒛_𝑖_ and 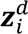 of each particle remains to be inferred. We adopt the variational autoencoder (VAE) architecture where the distributions of latent codes 𝒛 and 𝒛^*d*^ are inferred by an encoder network 𝑓. Specifically, OPUS-TOMO leverages a combination of the training objective of 𝛽-VAE^36^ and the structural disentanglement prior proposed in OPUS-DSD^29^. Let the dimension of 3D volume be *N*^3^, the training objective of OPUS-TOMO is of the form,

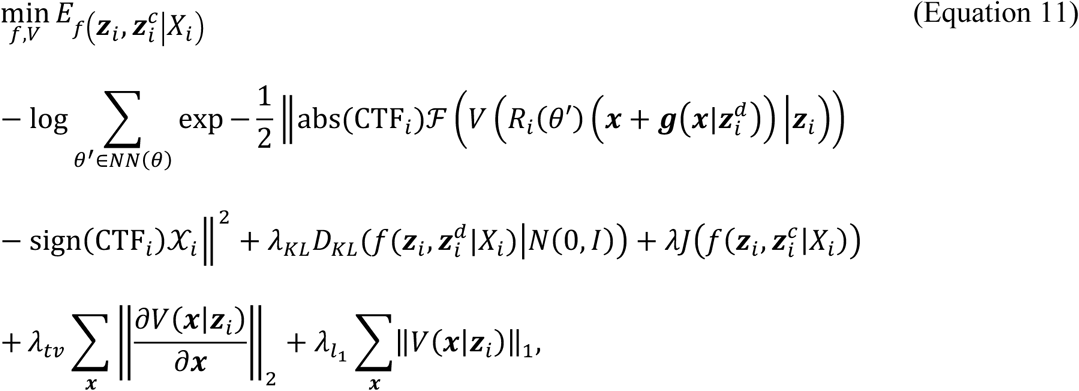

where the first term is expectation of error between the experimental subtomogram 𝑋_𝑖_ and the reconstruction over the distributions of latent codes and the nearest neighboring poses *NN*(𝜃) of 𝜃 in 𝑆𝑂(3), *N*(0, 𝐼) is the standard Gaussian distribution, 𝐷_𝐾𝐿_ is the KL divergence between the distributions of latent codes estimated by the encoder network 𝑓 and the standard Gaussian distribution, 𝜆_𝐾𝐿_ is the restraint strength for the KL divergence, 𝜆 is the restraint strength of structural disentanglement prior for the latent code of structure 𝒛, 𝜆_𝑡𝑣_ is the restraint strength for the total variation of structure, and 𝜆_𝑙1_ represents the restraint strength of the 𝐿_1_ norm of the structure. It is worth noting that the first expectation term in (Equation 11) is intractable, and approximated by the reparameterization trick during training ^34^.

### Experimental Protocol

We followed the conventional approaches to prepare subtomograms and performed subtomogram averaging before the structural heterogeneity analysis by OPUS-TOMO^13^. Due to inaccuracies in image tracking during data collection, the tilt series was first aligned to make 𝑦 axis as the tilt axis, and the center of image being the center of tilting before tomogram reconstruction. The alignment of tilt series can be performed either by AreTOMO^46^ or using the deposited parameters. The tomogram can then be reconstructed in WARP^6^ by importing the alignment parameters.

Template matching from pyTOM^16^ was used to select subtomograms from target species. For each tilt series, the subtomograms were sorted according to their correlation coefficients with respect to template calculated by pyTOM, and the subtomograms with correlation coefficients that were higher than a threshold was selected. We used WARP to export 3D subtomograms picked by template matching and the per-particle CTF parameters for them. With subtomograms and their 3D CTF parameters exported, we performed structural heterogeneity analysis by training OPUS-TOMO using the subtomograms with their poses from template matching. We select subtomgram clusters for target species by clustering the composition latent space learned by OPUS-TOMO using KMeans algorithm. Selected subtomogram sets with orientations determined by template matching were imported in M^4^ for high-resolution refinement.

### Training

The training process of OPUS-TOMO is similar to OPUS-DSD^29^. The subtomogram undergoes data augmentation processes before being supplied into the encoder. The data augmentation consists of several steps. Firstly, the Fourier transform of the subtomogram is blurred by a randomly sampled B-factor in the range [−0.25, 0.75] × 4𝜋^2^. The contrast and brightness of the subtomogram is also randomly altered according to the SNR of the subtomogram. Next, the subtomogram is multiplied with a randomly chosen constant in the range 1 + [−0.15,0.15] × SNR to change its overall contrast, and the subtomogram is added by a randomly chosen constant in the range [−0.2,0.2] × SNR to change its overall brightness. The encoder takes the randomly augmented subtomogram and outputs its latent codes. The decoders take the latent codes to reconstruct the corresponding 3D volume and deformation field for the input subtomogram. OPUS-TOMO reads the CTF parameters for the input subtomogram from STAR file and computes the corresponding 3D CTF. The decoders can then reconstruct the subtomogram using the 3D volume, deformation field, pose, and 3D CTF according the subtomogram formation model in (Equation 8). The reconstructed subtomogram and the original subtomogram for input together forms the reconstruction loss. We construct the training objective by combining the reconstruction loss, the priors for latent codes, and the total variation and 𝐿_1_ norms^52^ for the template volumes with specified weights as it is given in (Equation 11).

During training, OPUS-TOMO uses the same batching process as in OPUS-DSD^29^ to sample subtomograms with similar projection directions as a batch. OPUS-TOMO is fast to train. It only takes only 10 minutes to train 25,900 subtomograms of size 128x128x128 on 4 NVIDIA V100 GPUs for one epoch. This is much faster than the traditional 3D classification methods in RELION using the same hardware for the same number of data when the number of class is set to 4, which takes 1 hour for one iteration.

### Hyperparameters’ Settings

The training losses of OPUS-TOMO were minimized by the Adam optimizer^53^. The learning rate of Adam was set to 5 × 10^−5^. Let the SNR of dataset be SNR, the restraint strength 𝜆 for structural disentanglement prior was set to 0.5 × SNR, and the restraint strength 𝜆_𝐾𝐿_ for KL divergence between latent codes and standard Gaussian was set to 0.5 × SNR. The SNR was estimated *ad hoc* during training by treating the reconstruction from OPUS-TOMO as signal and the input subtomogram with reconstruction subtracted as noise. Formally, using a batch of *N* images, the SNR was estimated as,

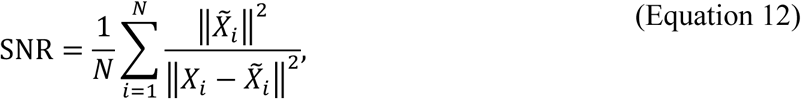

where *X̄*_𝑖_ was the subtomogram reconstructed by OPUS-TOMO, and 𝑋_𝑖_ was the experimental subtomogram. 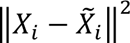 estimated the square of the norm of noises in the experimental subtomogram, while 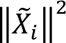 estimated the square of the norm of signals. The restraint strength 𝜆_𝑡𝑣_ for the total variation of 3D reconstruction was set to 0.1, and the restraint strength 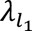 for the 𝐿_1_ norm of 3D reconstruction was set to 0.3.

## Data availability

We used the following publicly available datasets: EMPIAR-11830^37^ (cryo-FIB-milled lamellae of *C. reinhardtii*), EMPIAR-10988^39^ (cryo-FIB-milled lamellae of wild-type *S. pombe*), EMPIAR-10304^54^(*E. coli* 70S Ribosome), EMPIAR-10499^45^ (Cm-treated *M. pneumoniae*) and EMPIAR-11843^30^ (Cm-treated *M. pneumoniae*). The trained weights and structural heterogeneity analysis results are available at Zenodo at https://doi.org/10.5281/zenodo.12631920 ^55^.

## Code availability

The source code of OPUS-TOMO is available at https://github.com/alncat/opusTOMO, and also at Zenodo at https://doi.org/10.5281/zenodo.13626517 ^56^.

## Acknowledgements

J.M. wants to thank the support from the National Key Research and Development Program of China (No. 2024YFA1307502), the Science and Technology Innovation Plan of Shanghai Science and Technology Commission (No. 23JS1400200), and the Research Fund for International Senior Scientists (No. W2431060). The authors specially thank James Krieger for his contribution to the code improvement of OPUS-TOMO.

## Author Contributions

Z.L., Q.W. and J.M. conceived the work. Z.L. designed the algorithm and implemented the software. Z.L. and X.C. performed experiments and analyses. All authors wrote and approved the final manuscript.

## Extended Data Figure Legends

**Extended Data Figure 1.**
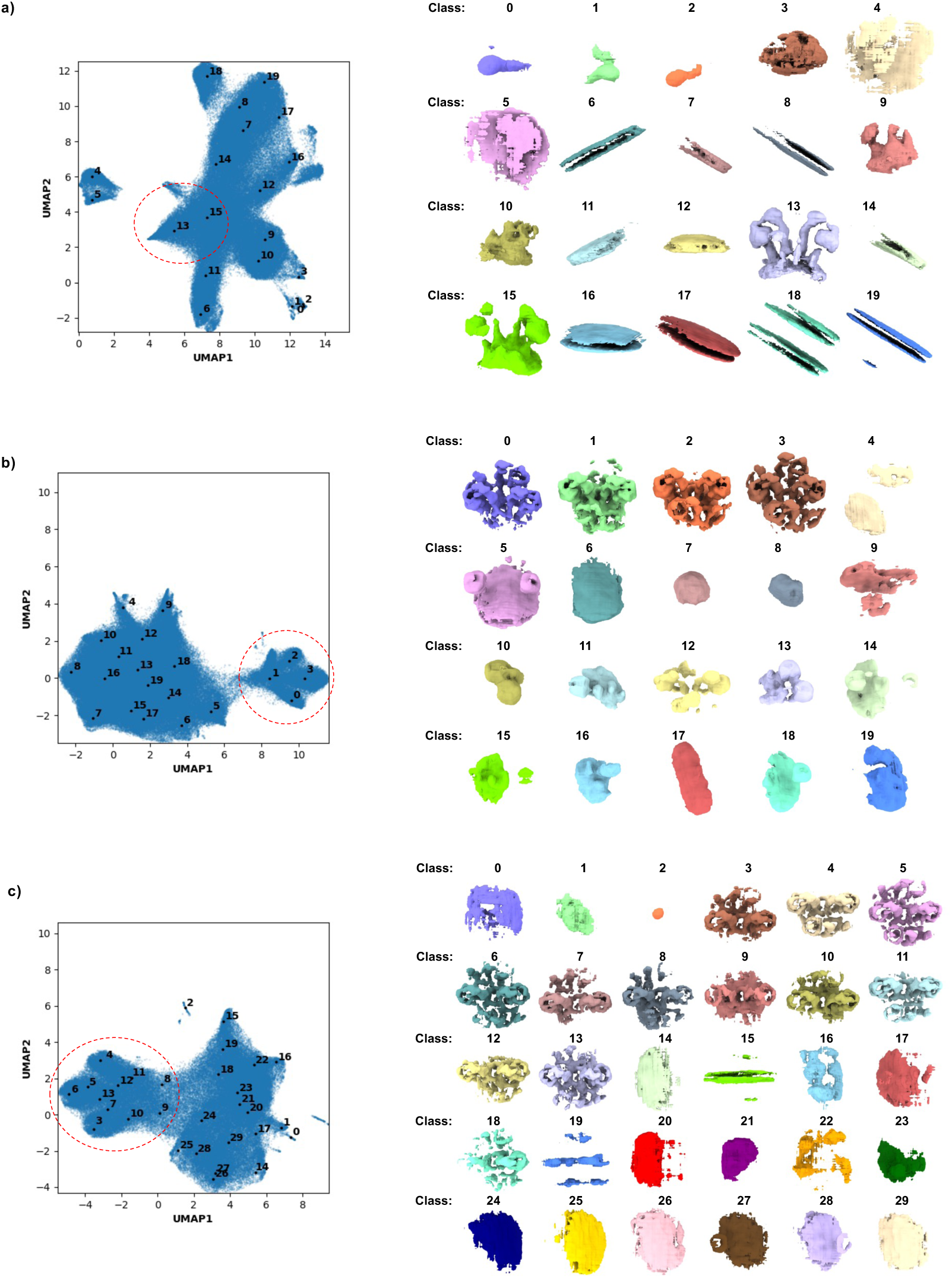
Heterogeneity analysis on the *C. reinhardtii* ATP synthase dimer (data from EMPIAR-11830). **a)**. (Left) UMAP visualization of the 12-dimensional composition latent space learned by OPUS-TOMO on template matching result and the distribution of centroids (black dots with labels) for clusters found by KMeans algorithm. (Right) Density maps at class centers reconstructed by the composition decoder of OPUS-TOMO. The cluster of classes showing densities for ATP synthase are marked by red dashed ellipse. **b)**. (Left) UMAP visualization of the 12-dimensional composition latent space learned by OPUS-TOMO on template matching result filtered by model trained in **a)** and the distribution of centroids (black dots with labels) for clusters found by KMeans algorithm. (Right) Density maps at class centers reconstructed by the composition decoder of OPUS-TOMO. **c)**. (Left) UMAP visualization of the 12-dimensional composition latent space learned by OPUS-TOMO on expert annotations and the distribution of centroids (black dots with labels) for clusters found by KMeans algorithm. (Right) Density maps at class centers reconstructed by the composition decoder of OPUS-TOMO.

**Extended Data Figure 2.**
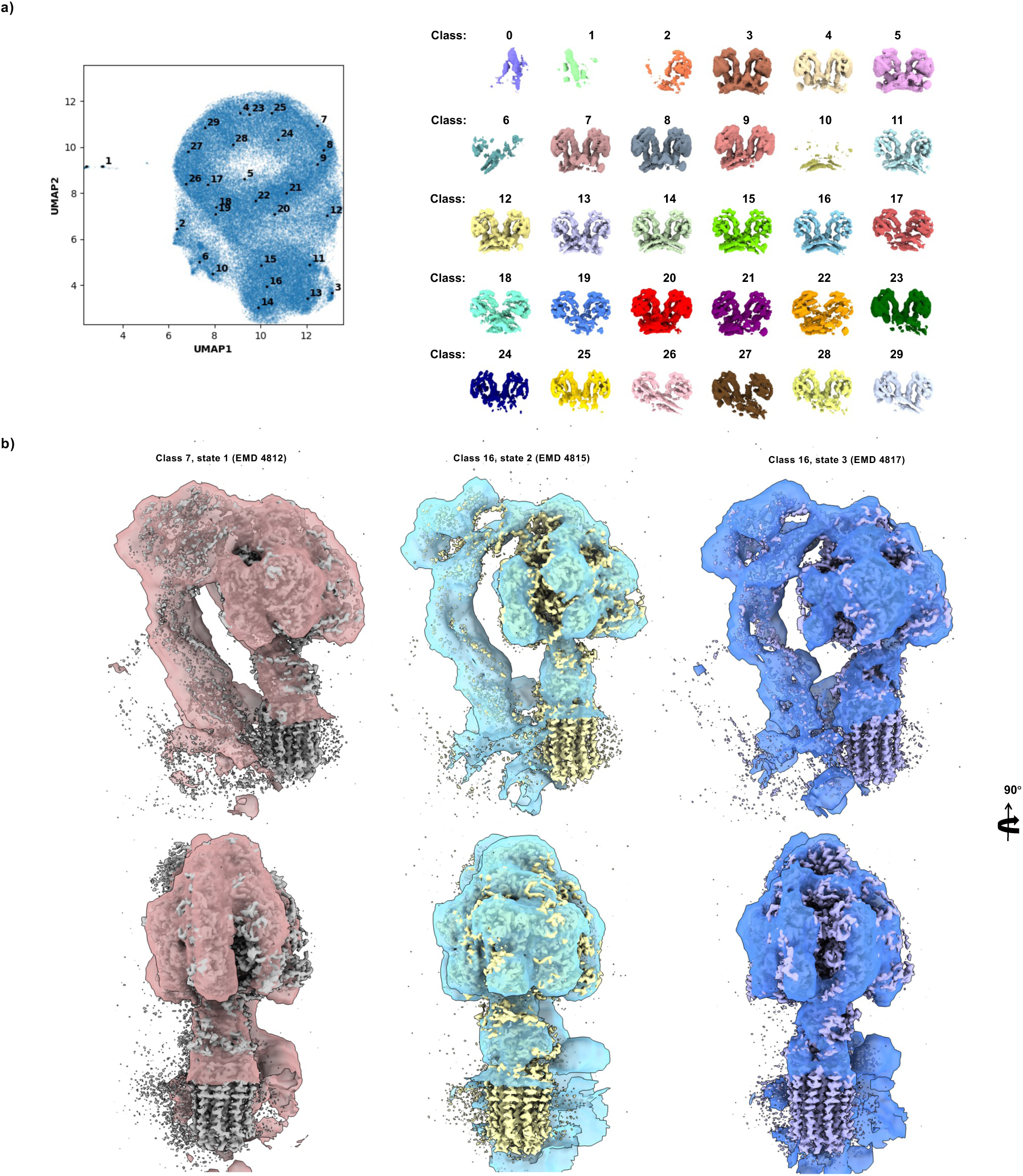
Heterogeneity analysis on the *C. reinhardtii* ATP synthase using M-refined expanded set. **a)**. (Left) UMAP visualization of the 12-dimensional composition latent space learned by OPUS-TOMO on M-refined expanded set and the distribution of centroids (black dots with labels) for clusters found by KMeans algorithm. (Right) Density maps at class centers reconstructed by the composition decoder of OPUS-TOMO. **b)**. Density maps of ATP synthase monomer from classes in **a)** overlaid with high-resolution density maps for state 1 (grey), state 2 (yellow), and state 3 (purple), (EMDB codes in parentheses).

**Extended Data Figure 3.**
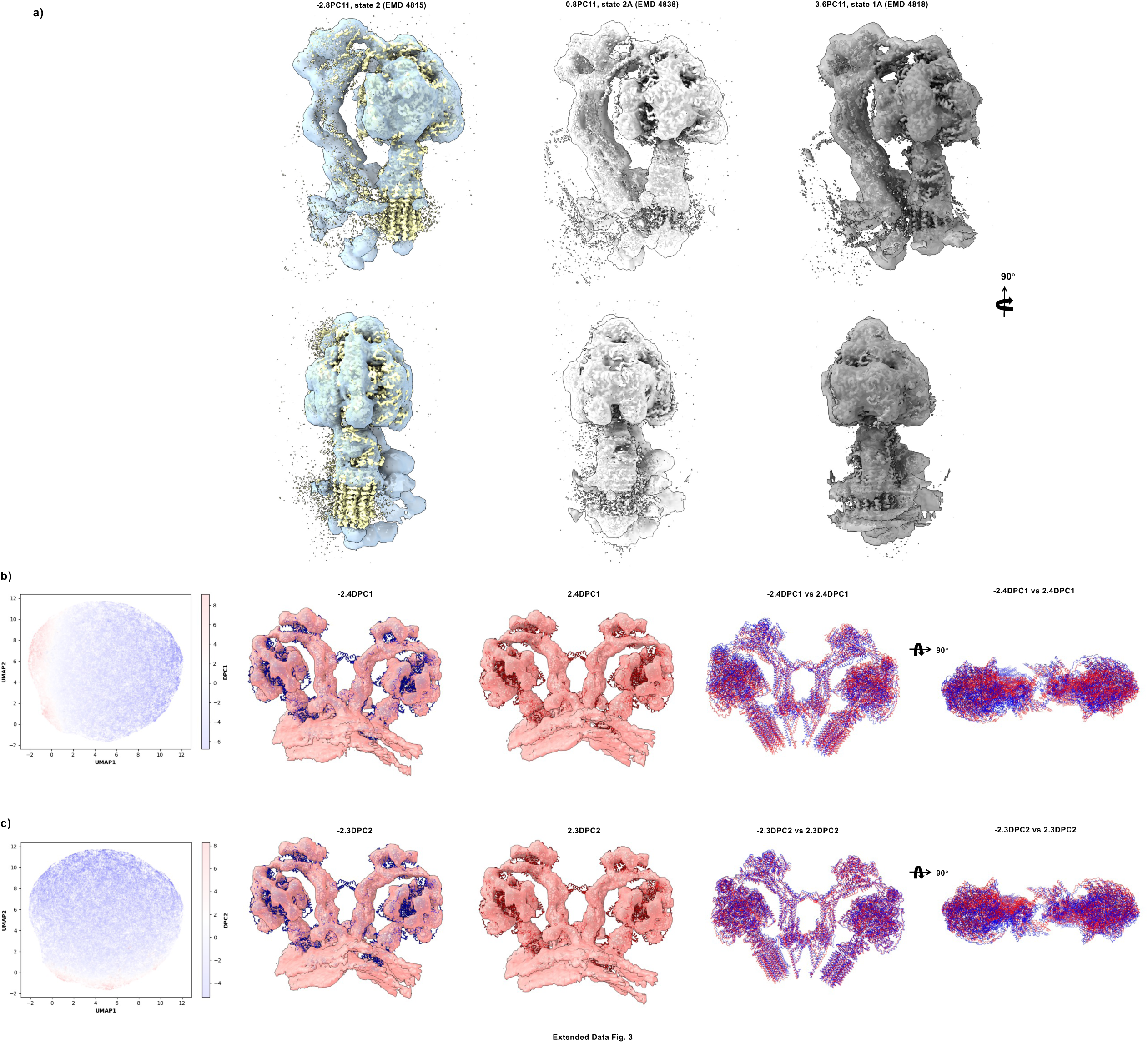
Rotary substates and dynamics of the *C. reinhardtii* ATP synthase reconstructed by OPUS-TOMO. **a)**. Density maps at different locations along the PC11 of composition latent space overlaid with high-resolution density maps for the corresponding rotary substate. **b)**. (Left) UMAP visualization of the dynamics latent space with latent codes colored by their locations an DPC1. The latent code in the positive end of DPC1 is colored in red, and the latent code in the negative end of DPC1 is colored in blue. The color scheme is consistent throughout all figures for UMAP visualization of dynamics latent space. (Middle) Density maps reconstructed by the dynamics decoder of OPUS-TOMO overlaid with their fitted structures. The structures of ATP synthase monomer (6RDE combined with peripheral stalk from 6RD4) are fitted separately. The structure fitted to density maps at negative end of DPC1 is colored in blue, while the structure fitted to density map at positive end of DCP1 is colored in red. (Right) Superposition of structures fitted to density maps at two ends of DPC1 reveals the relative displacements of residues when traversing along DPC1. **c)**. (Left) UMAP visualization of the dynamics latent space with latent codes colored by their locations on DPC2. The direction of DPC2 is orthogonal to DPC1. (Middle) Density maps reconstructed by the dynamics decoder of OPUS-TOMO overlaid with their fitted structures. (Right) Superposition of structures fitted to density maps at two ends of DPC2 reveals the relative displacements of residues when traversing along DPC2.

**Extended Data Figure 4.**
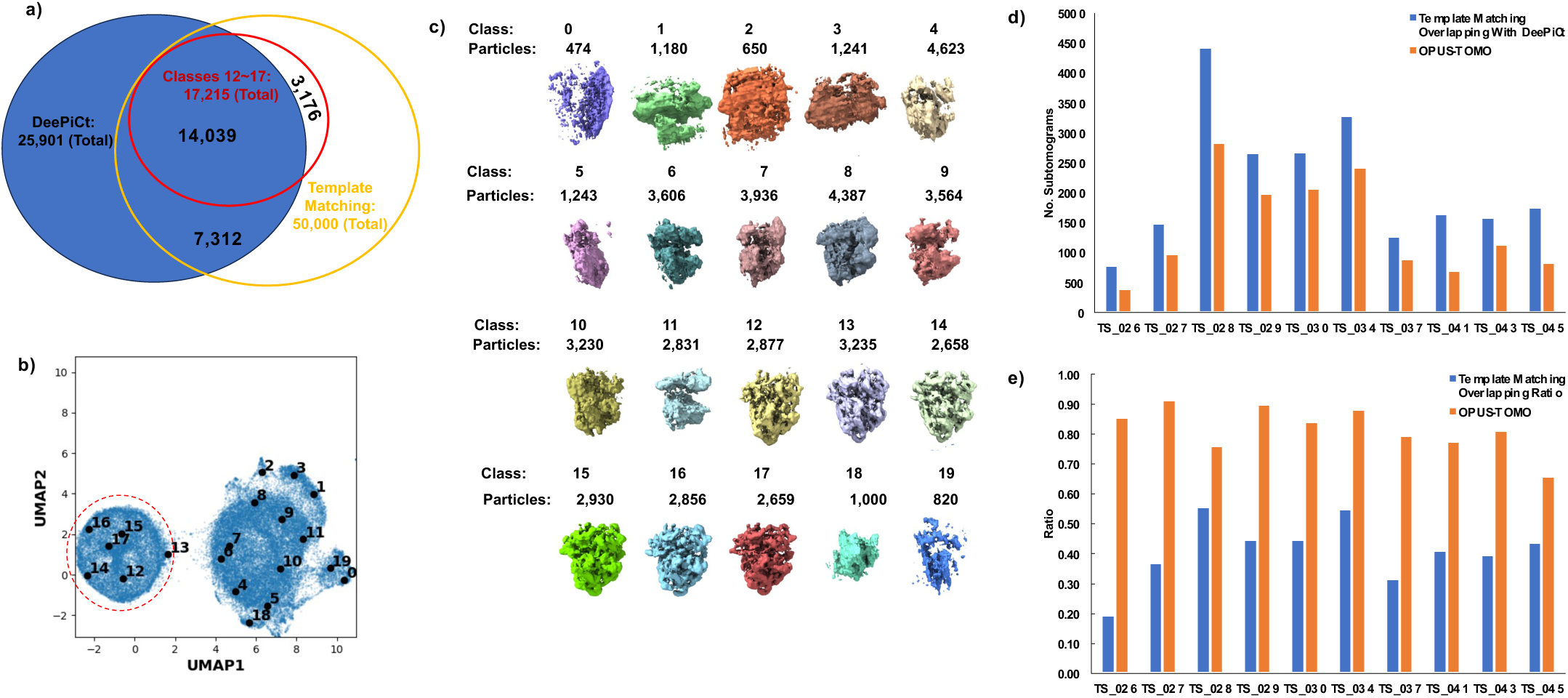
Purifying template matching result for the *S. pombe* 80S ribosome (data from EMPIAR-10988). **a).** Venn diagram for the subtomograms for 80S ribosome from different methods. The blue ellipse represents the subtomograms from DeePiCt’s expert annotation^39^ which is 25,901 in total. DeePiCt’s expert annotation is referred to as DeePiCt for simplicity. The yellow ellipse represents the subtomograms from our template matching, which is 50,000 in total. The red ellipse represents the cluster formed by Classes 12∼17 from OPUS-TOMO’s clustering result, which is 17,215 in total. The number of subtomograms in the intersection between DeePiCt and Classes 12∼17 is 14,039. The number of subtomograms in the intersection between DeePiCt and template matching is 21,351. **b)**. UMAP visualization of the 12-dimensional latent space learned by OPUS-TOMO and the distribution of centers for clusters found by KMeans algorithm. Solid black dot represents the cluster center for labelled class. Red dashed ellipse highlights the cluster of subtomograms corresponding to 80S ribosome. **c)**. Twenty density maps reconstructed by the composition decoder of OPUS-TOMO at centroids found by KMeans clustering. **d).** Distribution of the number of subtomograms from Classes 12∼17 given by OPUS-TOMO in each tilt series which overlap with DeePiCt’s result. “Template Matching Overlapping with DeePiCt” refers to the number of subtomograms from the template matching result that overlap with DeePiCt’s result in each tilt series. “OPUS-TOMO” refers to the number of subtomograms from Classes 12∼17 in OPUS-TOMO’s result that overlap with DeePiCt’s result in each tilt series. **e).** Distribution of the ratio of subtomograms from Class 12∼17 given by OPUS-TOMO in each tilt series which overlap with DeePiCt’s result. The ratio is calculated by dividing the number of subtomograms overlapping with DeePiCt’s result by the total number of subtomograms of the particle set for each tilt series. “Template Matching Overlapping Ratio” refers to the ratio of subtomograms in the template matching result that overlap with DeePiCt’s result in each tilt series. “OPUS-TOMO” refers to the ratio of subtomograms from Classes 12∼17 in OPUS-TOMO’s result that overlap with DeePiCt’s result in each tilt series.

**Extended Data Figure 5.**
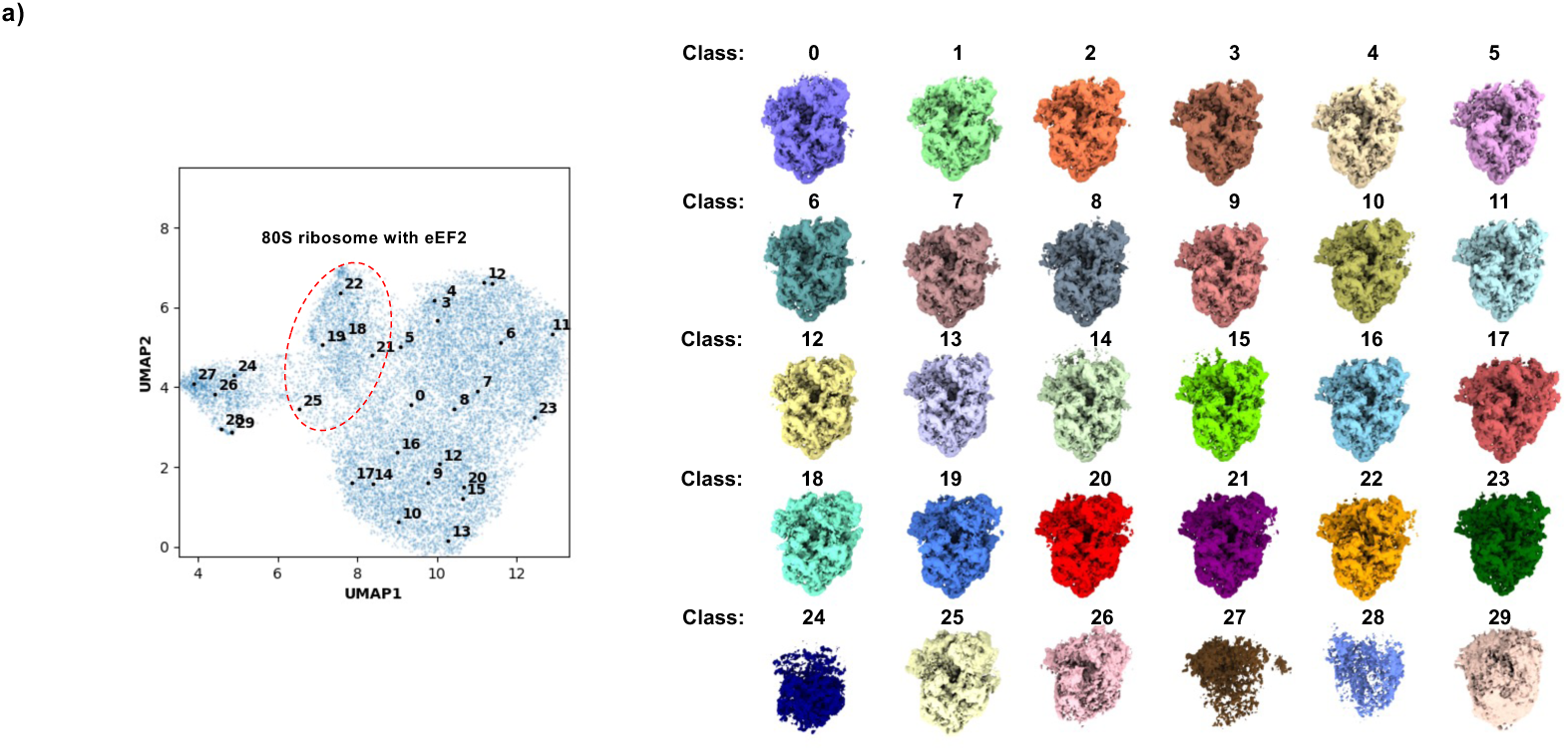
Heterogeneity analysis on the the *S. pombe* 80S ribosome (data from EMPIAR-10988). **a)**. (Left) UMAP visualization of the 12-dimensional composition latent space learned by OPUS-TOMO on M-refined subtomogram set purified by OPUS-TOMO and the distribution of centroids (black dots with labels) for clusters found by KMeans algorithm. (Right) Density maps at class centers reconstructed by the composition decoder of OPUS-TOMO. The cluster of classes showing 80S ribosome with eEF2 are marked by red dashed ellipse.

**Extended Data Figure 6.**
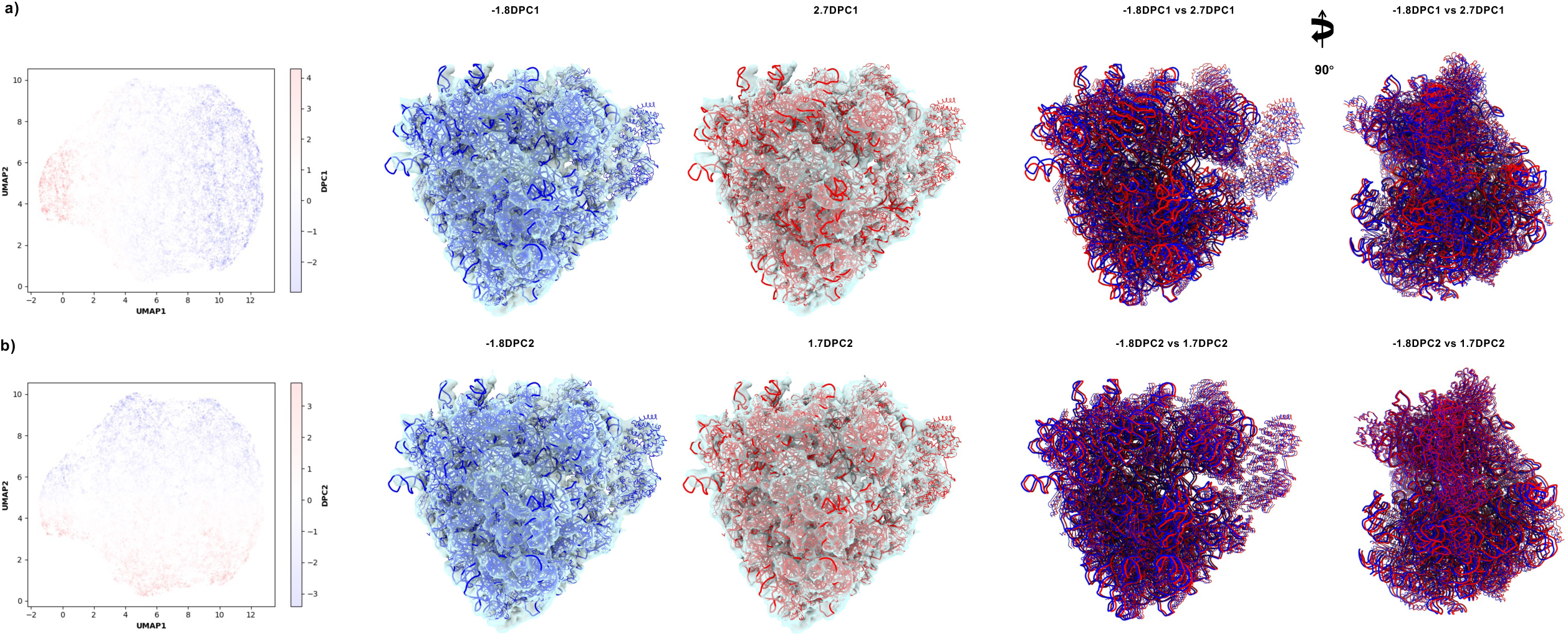
Inter-subunit dynamics of the *S. pombe* 80S ribosome reconstructed by the dynamics decoder of OPUS-TOMO. **a).** (Left) UMAP visualization of the dynamics latent space with latent codes colored by their locations on DPC1. (Middle) Density maps reconstructed by the dynamics decoder of OPUS-TOMO overlaid with their fitted structures. The structures of 40S and 60S subunit are fitted separately. The structure of 80S ribosome (PDB code: 7B7D) fitted to density maps at negative end of DPC1 is colored in blue, while the structure fitted to density map at positive end of DCP1 is colored in red. (Right) Superposition of structures fitted to density maps at two ends of DPC1 reveals the relative displacements of residues when traversing along DPC1. **b)**. (Left) UMAP visualization of the dynamics latent space with latent codes colored by their locations on DPC2. The direction of DPC2 is orthogonal to DPC1. (Middle) Density maps reconstructed by the dynamics decoder of OPUS-TOMO overlaid with their fitted structures. (Right) Superposition of structures fitted to density maps at two ends of DPC2 reveals the relative displacements of residues when traversing along DPC2.

**Extended Data Figure 7.**
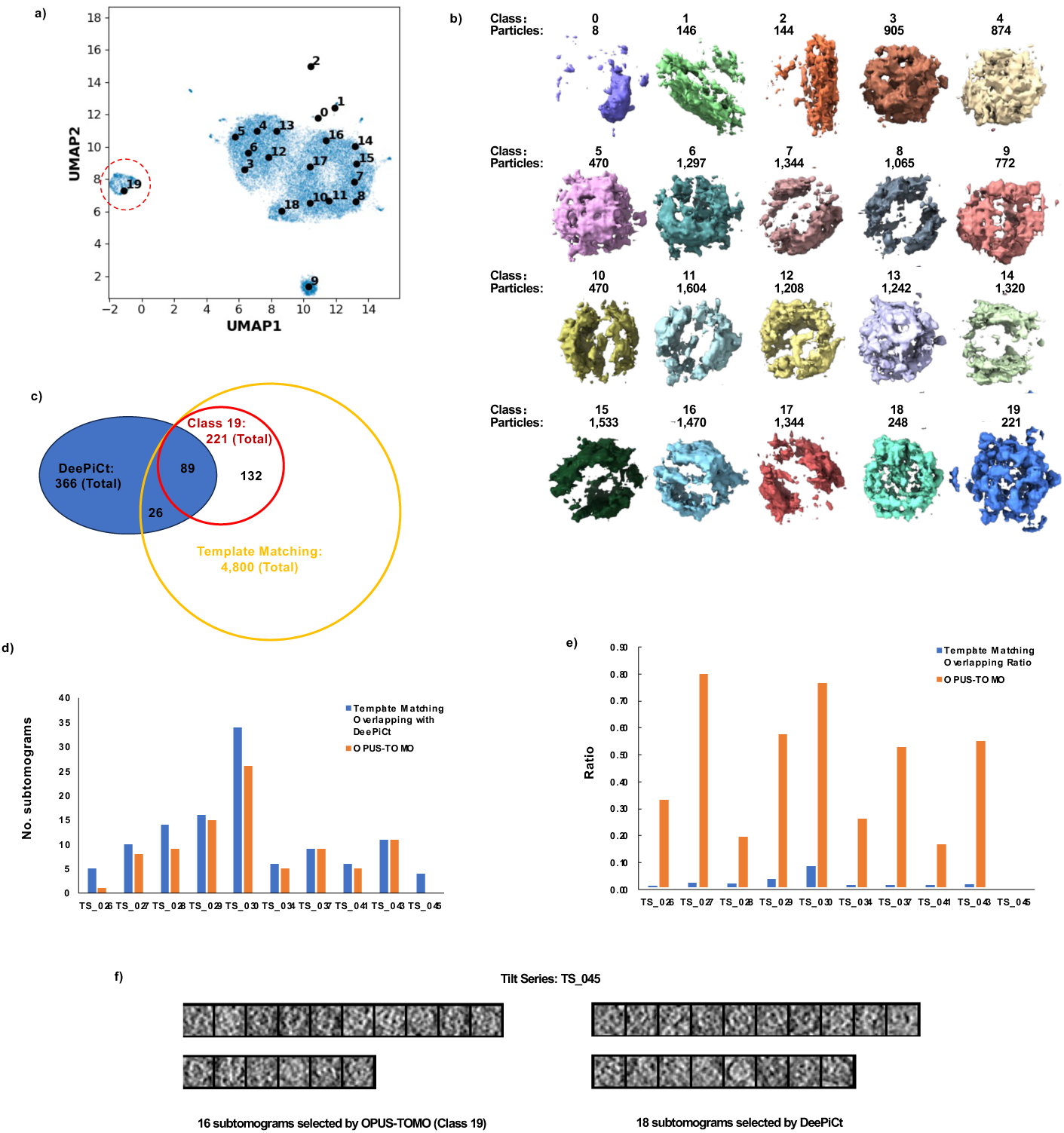
Heterogeneity analysis on the *S. pombe* FAS (EMPIAR-10988). **a)**. UMAP visualization of the 12-dimensional latent space learned by OPUS-TOMO and the distribution of centroids for clusters found by KMeans algorithm. Solid black dot represents the cluster center for labelled class. **b)**. Density maps reconstructed by the compostion decoder of OPUS-TOMO using twenty cluster centers as inputs. Class 19 show complete densities for FAS (red dashed ellipse in **a)**). **c)**. Venn diagram for the subtomograms for FAS from different methods. The blue ellipse represents the subtomograms from DeePiCt’s expert annotation^39^, which is 366 in total. DeePiCt’s expert annotation is referred to as DeePiCt for simplicity. The yellow ellipse represents the subtomograms from template matching^7^, which is 4,800 in total. The red ellipse represents the Class 19 from OPUS-TOMO’s clustering result, which is 221 in total. The number of subtomograms in the intersection between DeePiCt and Class 19 is 89. The number of subtomograms in the intersection between DeePiCt and template matching is 115. **d)**. Distribution of the number of subtomograms from Class 19 given by OPUS-TOMO in each tilt series which overlap with DeePiCt’s result. “Template Matching Overlapping with DeePiCt” refers to the number of subtomograms from the template matching result that overlap with DeePiCt’s result in each tilt series. “OPUS-TOMO” refers to the number of subtomograms from Class 19 in OPUS-TOMO’s result that overlap with DeePiCt’s result in each tilt series. **e)**. Distribution of the ratio of subtomograms from Class 19 given by OPUS-TOMO in each tilt series which overlap with DeePiCt’s result. The ratio is calculated by dividing the number of subtomograms overlapping with DeePiCt’s result by the total number of subtomograms of the particle set for each tilt series. “Template Matching Overlapping Ratio” refers to the ratio of subtomograms in the template matching result that overlap with DeePiCt’s result in each tilt series. “OPUS-TOMO” refers to the ratio of subtomograms from Class 19 in OPUS-TOMO’s result that overlap with DeePiCt’s result in each tilt series. **f)**. Projections of subtomograms from the tilt series TS_045 for Class 19 in OPUS-TOMO, and the projections of subtomograms from DeePiCt. The subtomograms are 4x binned and blurred by a Gaussian with standard deviation 1 before projecting along the 𝑍 axis.

**Extended Data Figure 8.**
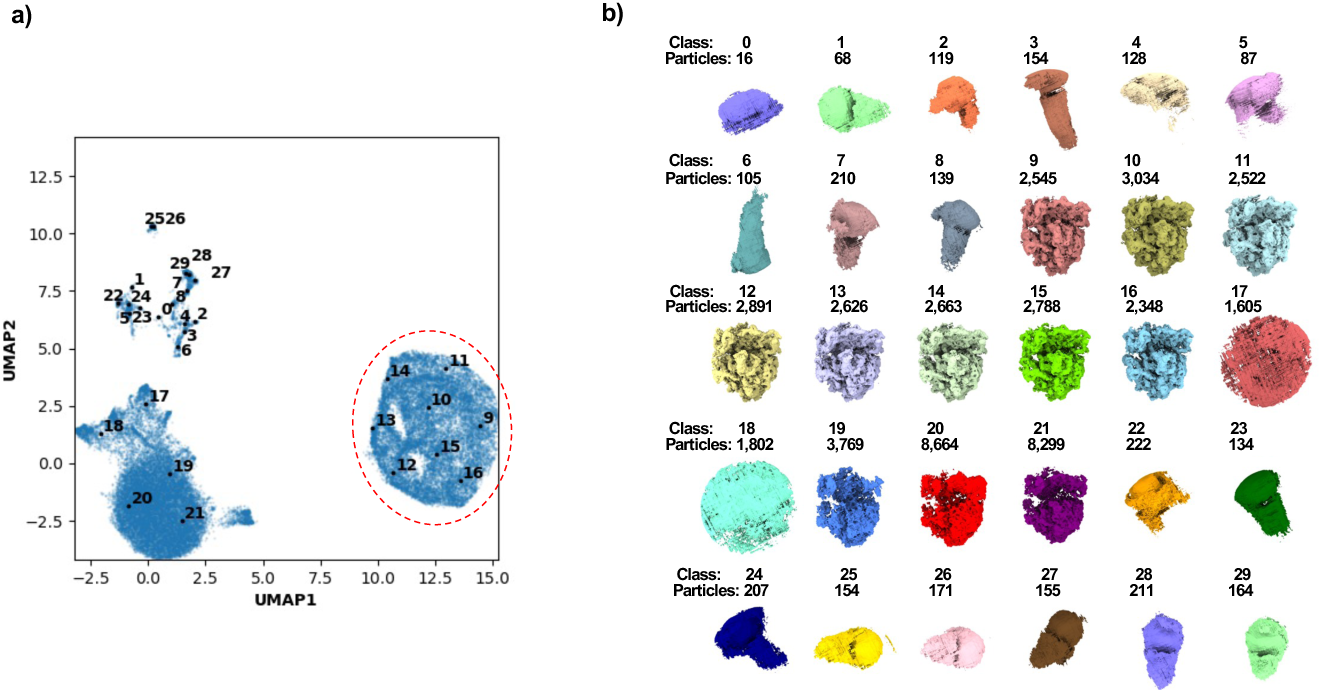
Purifying Template Matching Result for the *M. pneumoniae* 70S Ribosome (data from EMPIAR-10499). **a)**. UMAP visualization of the 12-dimensional composition latent space learned by OPUS-TOMO and the distribution of centroids for clusters found by KMeans algorithm. Solid black dot represents the cluster center for labelled class. Red dashed ellipse highlights the cluster of subtomograms corresponding to 70S ribosome. **b).** Thirty density maps at class centers reconstructed by the composition decoder of OPUS-TOMO.

**Extended Data Figure 9.**
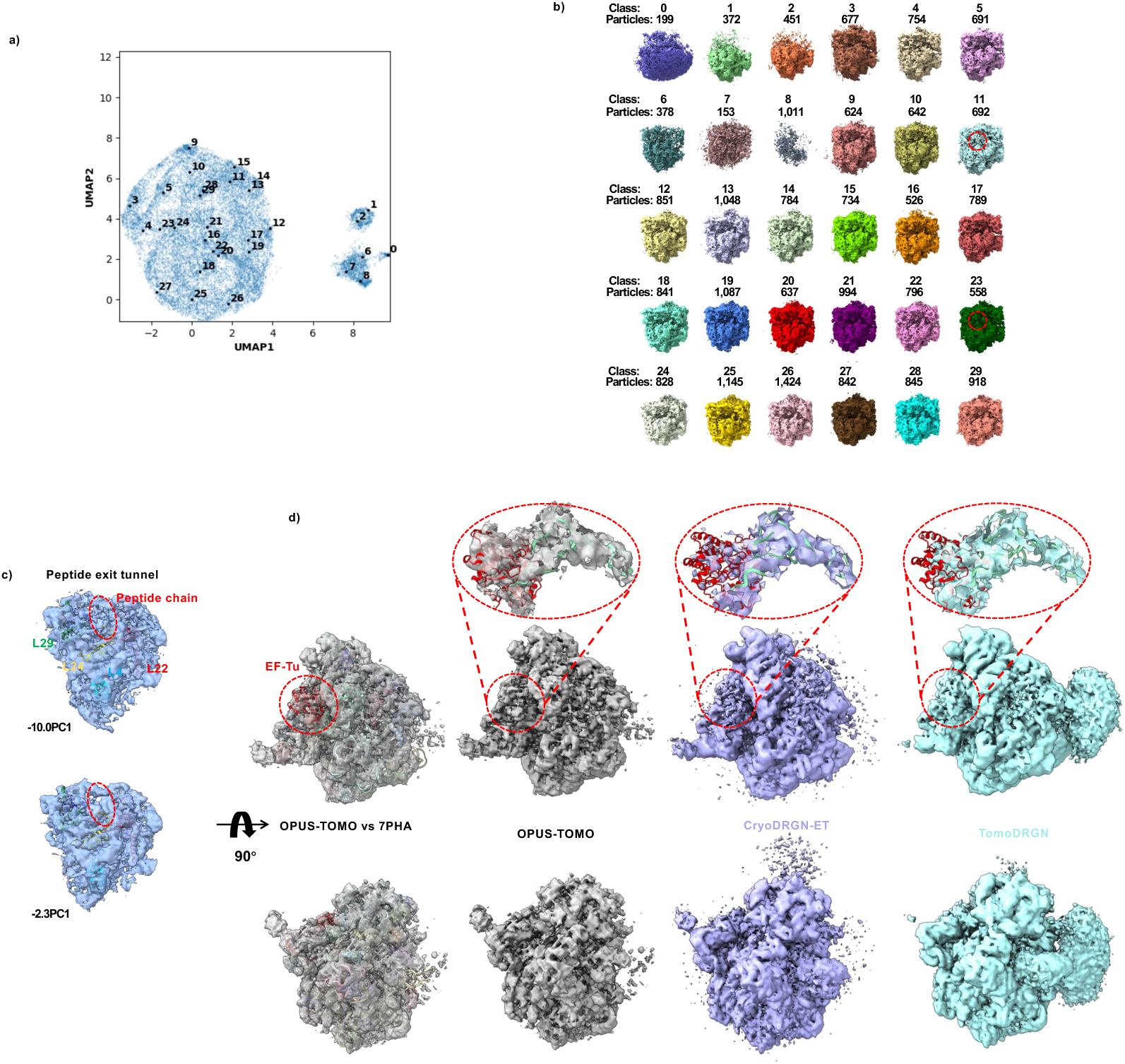
Heterogeneity analysis on the *M. pneumoniae* 70S ribosome (data from EMPIAR-11843). **a)**. UMAP visualization of the 12-dimensional latent space learned by OPUS-TOMO and the distribution of centroids for clusters found by KMeans algorithm. Solid black dot represents the cluster center for labelled class. **b)**. Thirty density maps at class centers reconstructed by the composition decoder of OPUS-TOMO. **c)**. The occupancy change at the peptide exit tunnel identified by OPUS-TOMO using latent codes at different locations along PC1. The locations of latent codes are specified as xPC1. The atomic model of 70S ribosome in Cm-treated *M. pneumoniae* cell (PDB accession code: 7PHA^49^) is fitted into the density map of − 10.0PC1. Red dashed ellipses mark the region at peptide exit tunnel. The subunits forming the peptide exit channel are colored and annotated by texts with the same colors. **d**). Comparison between 70S ribosome with EF-Tu cofactor reconstructed by different methods. The atomic model of 70S ribosome in Cm-treated *M. pneumoniae* cell (PDB accession code: 7PHA^49^) is fitted into the density map at 4.8PC9 reconstructed by OPUS-TOMO for reference. Red dashed ellipses mark the densities for EF-Tu-tRNA and their zoomed views. The atomic model of EF-Tu-tRNA from 7PHA is fitted into density maps and shown in zoomed views. The density maps are contoured by normalizing volumes of densities for EF-Tu-tRNA.

**Extended Data Figure 10.**
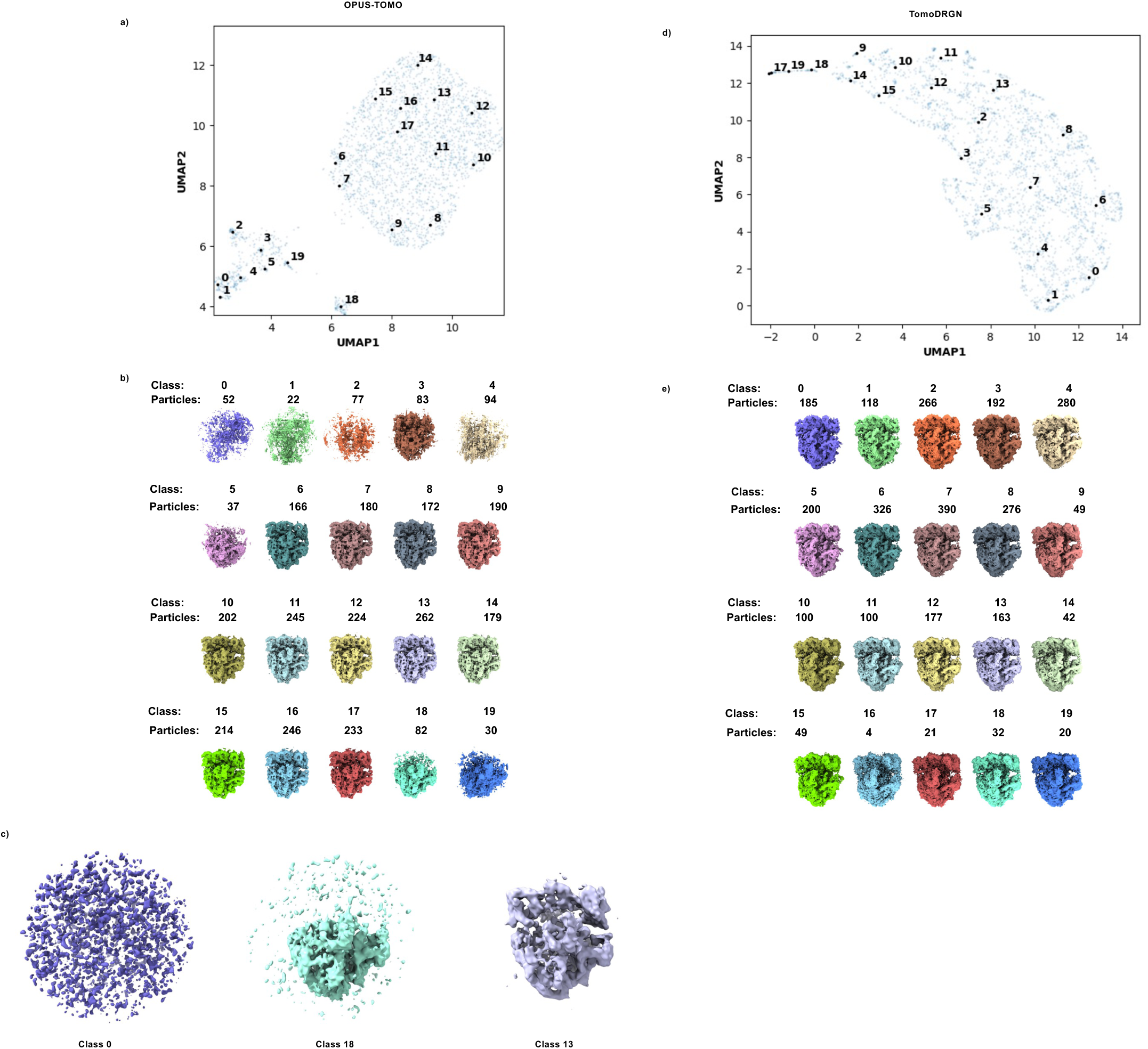
Comparison of the performances of heterogeneity analysis for different methods on the *M. pneumoniae* 70S ribosome (the first eight tomograms from EMPIAR-11843). **a).** UMAP visualization of the 12-dimensional latent space learned by OPUS-TOMO and the distribution of centroids for clusters found by KMeans algorithm. Solid black dot represents the cluster center for labelled class. **b)**. Twenty density maps at class centers reconstructed by the composition decoder of OPUS-TOMO. **c)**. Subtomogram averages for representative classes reconstructed by *relion_reconstruct* in RELION 3.0.8 using subtomograms from corresponding classes. The subtomogram averages are blurred by Gaussian kernel with standard deviation of 2. **d)**. UMAP visualization of the 8-dimensional latent space learned by tomoDRGN and the distribution of centroids for clusters found by KMeans algorithm. Solid black dot represents the cluster center for labelled class. **e)**. Twenty density maps at class centers reconstructed by the decoder of tomoDRGN.

## Extended Data Table

**Extended Data Table 1.**
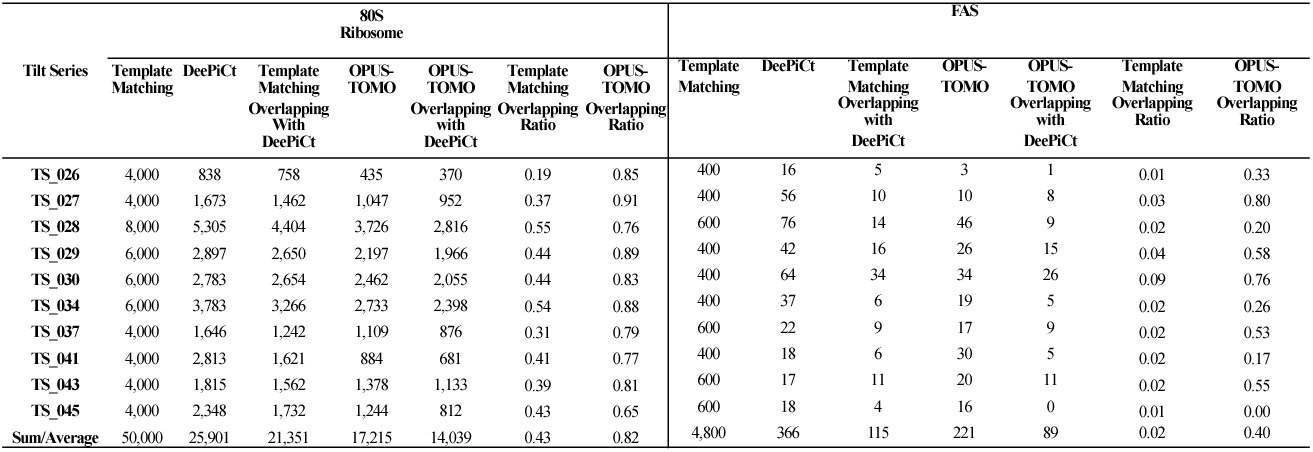
Distributions of subtomograms representing 80S ribosome (Left) or FAS (Right) in each tilt series. “Template matching” refers to the number of subtomograms localized by template matching; “DeePiCt” refers to the number of subtomograms which are extracted according to expert annotations from DeePiCt^39^; “Template matching overlapping with DeePiCt” refers to the number of subtomograms that in both template matching and DeePiCt’s annotation result; “OPUS-TOMO” refers to the number of subtomograms from Classes 12∼17 (for 80S ribosome, **Extended Data Fig. 4c∼d**) or Class 19 (for FAS, **Extended Data Fig. 7a∼b**) given by OPUS-TOMO in each tilt series; and “OPUS-TOMO Overlapping with DeePiCt” refers to the number of subtomograms that are in both Classes 12∼17 (for 80S ribosome) or Class 19 (for FAS) from OPUS-TOMO’s clustering result and DeePiCt’s annotation result. The ratio was calculated by the number of overlapping subtomograms divided by the number of subtomograms from template matching (for “Template matching overlapping ratio”) or from OPUS-TOMO’s clustering result (for “OPUS-TOMO overlapping ratio), respectively.

## Supplementary Video Legends

**Supplementary Video 1.** Traversal of PC11 of the composition latent space of the *C. reinhardtii* ATP synthase reveals that the central stalk rotates by 120° while the F_1_ Head rotates by 30° during the transition from state 2 to state 1A. The structure for ATP synthase in state 2 (PDB code: 6RDE) is shown for reference. The video is created by traversing PC11 forward and backward twice.

**Supplementary Video 2.** Traversal of DPC1 of the dynamics latent space of the *C. reinhardtii* ATP synthase reveals the global rotation of ATP synthase dimer. The video is created by traversing DPC1 forward and backward twice.

**Supplementary Video 3.** Traversal of DPC2 of the dynamics latent space of the *C. reinhardtii* ATP synthase reveals the twist of ATP synthase dimer resulting from the rotation of ATP synthase monomer in opposite direcitons. The video is created by traversing DPC2 forward and backward twice.

**Supplementary Video 4.** Traversal of PC3 of the composition latent space of the *S. pombe* 80S ribosome reveals the transition from the state with A/T- and P-site tRNAs to the state with classical A/P- and P/E-site tRNAs, facilitating by the L1 stalk. The L1 stalk moves to E-site and binds with P/E-site tRNA during transition. The 40S subunit rotates counter-clockwise. The atomic models for A/T-site tRNA (yellow), A-site tRNA (green), P-site tRNA (red) and E-site tRNA (blue) are shown for reference. The video is created by traversing PC3 forward and backward twice. The dynamics of tRNAs can be identified by observing the occupancy changes at the A-, P- and E-sites with atomic models.

**Supplementary Video 5.** Traversal of DPC1 of the dynamics latent space of the *S. pombe* 80S ribosome reveals that 40S and 60S subunits rotate in opposite direction. The video is created by traversing DPC1 forward and backward twice.

**Supplementary Video 6.** Traversal of DPC2 of the dynamics latent space of the *S. pombe* 80S ribosome reveals the 40S and 60S subunits rotate in the same direction while translating rightward. The video is created by traversing DPC2 forward and backward twice.

**Supplementary Video 7.** Traversal of PC9 of the composition latent space of the *M. pneumoniae* 70S ribosome reveals the transition from the state with EF-Tu-tRNA and P- and E-site tRNA to the state with A- and P- site tRNAs, corresponding to the delivery of tRNA into A-site and the exit of E-site tRNA. The atomic models for EF-Tu (blue), tRNA (cyan), A-site tRNA (yellow), P-site tRNA (red) and E-site tRNA (green) are shown for reference. The directions of movements during forward traversal are marked by red arrows. The video is created by traversing PC9 backward and forward twice with and without cropping. The dynamics of tRNAs can be identified by observing the occupancy changes at the A- and P- sites with atomic models. The dynamics of the subunits can be identified by observing the density changes in regions indicating by red arrows.

**Supplementary Video 8.** Translocation dynamics reconstructed by cryoDRGN-ET and tomoDRGN. The heterogeneity analyses of cryoDRGN-ET and tomoDRGN are performed using multi-particle refinement result from M at a resolution of ∼3.5 Å. The dynamics reconstructed by cryoDRGN-ET (**Left**) is created using the first 32 deposited volumes, which are reconstructed at latent codes sampled by interpolation between manually chosen states^31^. The dynamics reconstructed by tomoDRGN (**Right**) is created using deposited volumes for Classes 23∼32 from the 100 classes clustered by KMeans algorithm^30^. The atomic models for EF-Tu, tRNA, A-site tRNA, and P-site tRNA are colored in blue, cyan, yellow, and red, respectively. The head and body of the 30S subunit and the L1 stalk are indicated by corresponding labels. The dynamics of tRNAs can be identified by observing the occupancy changes at the A- and P-sites with atomic models. The dynamics of the subunits located between the head and body can be identified by observing the density changes indicating by red arrows.

